# Changes in a new type of genomic accordion may open the pallets to increased monkeypox transmissibility

**DOI:** 10.1101/2022.09.30.510261

**Authors:** Sara Monzón, Sarai Varona, Anabel Negredo, Juan Angel Patiño-Galindo, Santiago Vidal-Freire, Angel Zaballos, Eva Orviz, Oskar Ayerdi, Ana Muñoz-García, Alberto Delgado-Iribarren, Vicente Estrada, Cristina García, Francisca Molero, Patricia Sánchez, Montserrat Torres, Ana Vázquez, Juan-Carlos Galán, Ignacio Torres, Manuel Causse del Río, Laura Merino, Marcos López, Alicia Galar, Laura Cardeñoso, Almudena Gutiérrez, Cristina Loras, Isabel Escribano, Marta Elena Alvarez-Argüelles, Leticia del Río, María Simón, MªAngeles Meléndez, Juan Camacho, Laura Herrero, Pilar Jiménez Sancho, Maria Luisa Navarro-Rico, Jens H. Kuhn, Mariano Sanchez-Lockhart, Nicholas Di Paola, Jeffrey R. Kugelman, Elaina Giannetti, Susana Guerra, Adolfo García-Sastre, Gustavo Palacios, Isabel Cuesta, Maripaz P. Sánchez-Seco

## Abstract

The currently expanding monkeypox epidemic is caused by a subclade IIb descendant of a monkeypox virus (MPXV) lineage traced back to Nigeria in 1971. In contrast to monkeypox cases caused by clade I and subclade IIa MPXV, the prognosis of current cases is generally favorable, but person-to-person transmission is much more efficient. MPXV evolution is driven by selective pressure from hosts and loss of virus–host interacting genes. However, there is no satisfactory genetic explanation using single-nucleotide polymorphisms (SNPs) for the observed increased MPXV transmissibility. We hypothesized that key genomic changes may occur in the genome’s low-complexity regions (LCRs), which are highly challenging to sequence and have been dismissed as uninformative. Using a combination of highly sensitive techniques, we determined a first high-quality MPXV genome sequence of a representative of the current epidemic with LCRs resolved at unprecedented accuracy. This effort revealed significant variation in short-tandem repeats within LCRs. We demonstrate that LCR entropy in the MPXV genome is significantly higher than that of SNPs and that LCRs are not randomly distributed. *In silico* analyses indicate that expression, translation, stability, or function of MPXV orthologous poxvirus genes (OPGs) 153, 204, and 208 could be affected in a manner consistent with the established “genomic accordion” evolutionary strategies of orthopoxviruses. Consequently, we posit that genomic studies focusing on phenotypic MPXV clade-/subclade-/lineage-/strain differences should change their focus to the study of LCR variability instead of SNP variability.

## INTRODUCTION

Monkeypox virus (MPXV) is a double-stranded DNA virus that belongs to genus *Orthopoxvirus* (varidnavirian *Nucleocytoviricota*: *Poxviridae*: *Chordopoxvirinae*) along with other human viruses, such as vaccinia virus (VACV) and variola virus (VARV) (International Committee on Taxonomy of Viruses, 2022). First encountered in 1958 in crab-eating macaques imported to Belgium (Magnus et al., 2009), MPXV has caused sporadic disease outbreaks in humans since the 1970s in Eastern, Middle, and Western Africa, totaling approximately 25,000 cases (case fatality rate 1–10%) (Beer and Rao, 2019), and also sporadic disease outbreaks among wild monkeys and apes (Patrono et al., 2020; Radonić et al., 2014). Exposure to MPXV animal reservoirs, in particular rope squirrels and sun squirrels, is a significant risk factor of human infections (Khodakevich et al., 1988).

The human disease caused by MPXV is designated as monkeypox in the World Health Organization (WHO) International Classification of Diseases, Eleventh Revision (ICD-11; code 1E71) (World Health Organization, 2022a). Phylogenetically, historic MPXV isolates cluster into two clades (Likos et al., 2005), designated I and II (Happi et al., 2022; World Health Organization, 2022b). Clade I viruses are considered more virulent and transmissible than clade II viruses (Damon, 2011; Likos *et al*., 2005; World Health Organization, 2022b).

Since May 2022, multiple European countries have reported a continuously increasing number of MPXV infections and associated disease, including clusters of cases associated with potential superspreading events in Belgium, Spain, and the United Kingdom (UK). As of September 30, 2022, a total of 68,428 cases had been reported in 106 countries/territories/areas in all six WHO regions. Of these, 67,739 were in 99 countries that have not reported MPXV infections prior to 2022 (Centers for Disease Control and Prevention, 2022). This rapid increase in infections prompted WHO to declare this epidemic a Public Health Emergency of International Concern (PHEIC) (Nuzzo et al., 2022). The viruses of the 2022 epidemic belong to subclade IIb (Antinori et al., 2022; Nextstrain, 2022; Vivancos et al., 2022), a line of descent of MPXV that had been circulating in Nigeria, likely since 1971 (Faye et al., 2018).

The clinical presentation monkeypox caused by clade I or IIa includes fever, headache, lymphadenopathy, and/or malaise, followed by a characteristic rash that progresses centrifugally from maculopapules via vesicles and pustules to crusts that may occur on the face, body, mucous membranes, palms of the hands, and soles of the feet (Ježek et al., 1987). The clinical presentation of the current MPXV subclade IIb infections diverges from classical monkeypox by having a good prognosis, self-limiting but infectious skin lesions (typically emerging at and restricted to the genital, perineal/perianal, and/or peri-oral areas) before the development of fever, lymphadenopathy, and malaise. Generalized disease usually manifests with a rash that has not been widely observed in the current outbreak. Human-to-human transmission is substantially higher in subclade IIb MPXV-associated outbreaks than in clade I and clade IIa MPXV (Bunge et al., 2022; Otu et al., 2022; Thornhill et al., 2022; Ulaeto et al., 2022; Vusirikala et al., 2022). The R_0_ for MXPV IIb among men who have sex with men (MSM) is higher than 1. Transmission may be catalyzed by decreasing protection associated with the VARV/smallpox vaccination campaign that ended in 1980 (Grant et al., 2020; Rimoin et al., 2010). Furthermore, a change in transmission route may be the cause of the difference in clinical presentation and pathogenesis as was shown in animal models (Reynolds et al., 2006).

Orthopoxvirus infections are classified as systemic or localized illnesses. Localized usually means that signs are restricted to the site of MPXV entry, as described in the 2022 outbreak. The involved orthopoxvirus, its route of entry, and the immune status of the host are usually the only determinants of generalized or localized infection. Different mechanisms of virion entry and egress, as well as virus-encoded host factors, are the main biological determinants (Liu et al., 2019; McFadden, 2005; Moss, 2006; 2016; Roberts and Smith, 2008). Changes in the genome of the current MPXV variant, such as gene loss (Kugelman et al., 2014), may explain both trends.

The MPXV genome is a linear, ≈197-kb-long double-stranded DNA with covalently closed hairpin ends. The genome’s densely packed orthologous poxvirus genes (OPGs) (Senkevich et al., 2021) are distributed over a central conserved region (“core”) and flanking terminal regions, each of which ends in identical but oppositely oriented ≈6.4-kb-long terminal inverted repetitions (ITRs). Roughly 193 open reading frames (ORFs) encode proteins with ≥60 amino-acid residues. “Housekeeping” proteins involved in MPXV transcription, replication, and virion assembly are encoded by OPGs located in the central conserved region, whereas proteins involved in host range and pathogenesis are mostly encoded by OPGs located in the terminal regions. Like all orthopoxvirus genomes, the MPXV genome contains numerous tandem repeats in the ITRs as well as nucleotide homopolymers all over the genome (Moss and Smith, 2021; Shchelkunov et al., 2002; Wittek and Moss, 1980). However, we also observed other similar structures through the MPXV genome in the form of short tandem repeats (STRs). Moreover, initial observations appear to indicate that these STRs (which may consist of dinucleotide, trinucleotide, or more complex palindromic repeats) are localized in areas where more variation is observed suggesting a crucial role in the MXPV biology and evolution.

Orthopoxviruses rapidly acquire higher fitness by massive gene amplification (genome expansion) when encountering severe bottlenecks *in vitro*. This amplification, akin to gene reduplication in organismal evolution, enables gene copies to accumulate mutations, potentially resulting in new protein variants that can overcome the bottlenecks. Subsequent gene copy reduction (genome contraction) offsets the costs associated with increasing genome length, thereby retaining the adaptive mutations (Elde et al., 2012). Orthopoxviruses also rapidly adapt to selective pressures by single-nucleotide insertions (genome expansion) or deletions (genome contractions) within poly-A or poly-T stretches, resulting in easily reversible gene-inactivating or re-activating frameshifts (Senkevich et al., 2020). These rhythmic genome expansions and contractions are referred to as “genomic accordions” at the gene and base level (Elde *et al*., 2012). Given the overall conservation of STRs in orthopoxvirus genomes, we hypothesized that their variation could be a third type of genomic accordion and that, overall, this type of adaptation (which we designate here as low-complexity regions [LCRs]), rather than single-nucleotide polymorphisms (SNPs), could be the key to understanding the unusual epidemiology of 2022 subclade IIb MPXV.

## RESULTS

### *De novo* assembly of subclade II lineage B.1 monkeypox virus (MPXV) genome sequence 353R

Using a template-based mapping approach, shotgun metagenomic short-read-based sequencing of nucleic acids in vesicular lesion swabs from Spanish monkeypox patients resulted in the determination of 48 MPXV consensus genome sequences with at least 10X read depth. A median of 39,697,742 high-quality reads per swab (maximum = 111,030,976; minimum = 7,780,032) were obtained using a NovaSeq 6000 Illumina sequencer. Although 98.12% of the reads were assigned as being of human origin, a median of 74,085 MPXV reads (maximum = 27,516,891; minimum = 30,854) sufficed to cover >99% of the genome (**Table S1**).

Read mapping indicated that, as expected, LCRs of the MPXV genome were mostly unresolved. More importantly, those results were biased by the MPXV genome sequence used for mapping (subclade II lineage A MPXV isolate MPXV-M5312_HM12_Rivers): LCRs were resolved by the used reference-mapping software tools “following” the sequence provided in the reference genome instead of reporting the actual sequence (**Figure S1A**). To determine the actual LCR sequence, we explored different assembly strategies generally used for resolving eukaryotic genomes, which mostly combine different sequencing technologies. To increase the chances of success, we applied these technologies to a monkeypox patient sample with a high proportion of high-quality viral reads (swab 353R). The *de novo* assembly obtained from NovaSeq (2×150-bp pair-ended reads), MiSeq (2×300-bp pair-ended reads), and Nanopore sequencing generated 3, 2, and 1 contigs belonging to MPXV, covering 97%, 97%, and 101% of the MPXV-M5312_HM12_Rivers sequence, respectively (**Figure 1**).

**Figure 1.**
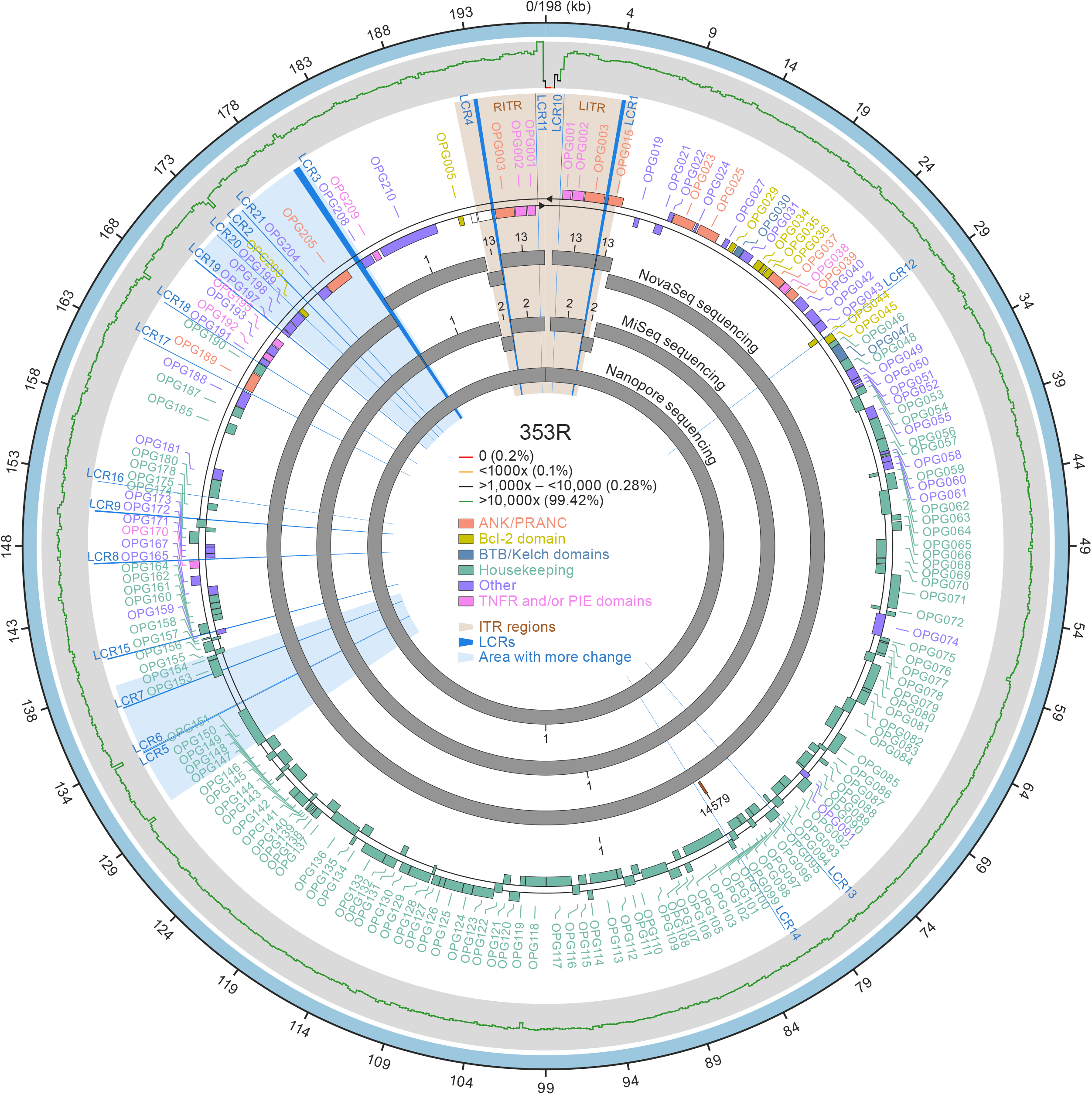
*De novo* assembly of subclade II lineage B.1 monkeypox virus (MPXV) genome sequence 353R. A visual representation of the fully annotated MPXV isolate 353R genome (based on the subclade II lineage A reference isolate MPXV-M5312_HM12_Rivers genome sequence annotation). Shown are (from the outside to the inside): high-quality genome (HQG) hybrid assembly (wide outer light blue ring); sequencing coverage distribution graph (thin ragged line [green: `≥`10,000x, 99.42%; black: 1,000x–10,000, 0.28%; orange: <1,000x-10, 0.1%; red: <10-0, 0.2%]); orthologous poxvirus gene (OPG) annotations according to the standardized nomenclature (Senkevich *et al*., 2021) (lettering and shaded boxes [orange: ANK/PRANC (N-terminal ankyrin protein with PRANC domain) inverted terminal repetition [ITR] regions; gold: Bcl-2 domain; blue: BTB/Kelch domains; green: housekeeping; purple: other; pink: TNFR and/or PIE domains); contigs from NovaSeq, MiSeq, and Nanopore sequencing (wide inner gray rings). Additionally, radial lines and shading that originate in the center and reach outward on the white background indicate low-complexity regions (LCRs; royal blue) and areas with more change (light blue).

### Characterization and validation of non-randomly distributed low-complexity regions (LCRs) in monkeypox virus (MPXV) genome sequence 353R

We applied a systematic approach for LCR discovery to the MPXV 353R sequence that resulted in the identification of 21 LCRs (13 STRs, 8 homopolymers; **Tables 1** and **S2**). Two pairs of LCRs (1/4 and 10/11) are located in the ITRs and are identical copies in reverse-complementary form. Consequently, we moved forward with 19 LCRs.

**Table 1.**
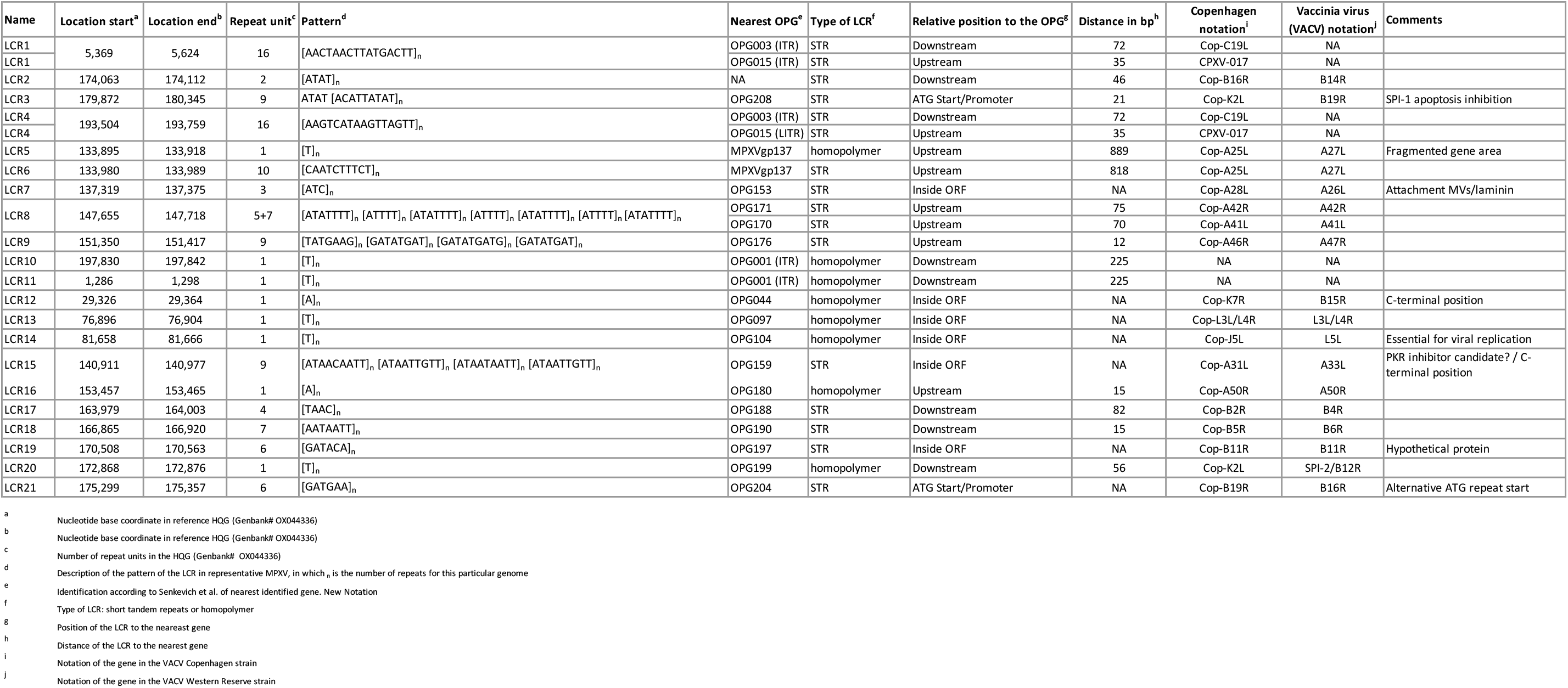
Low-complexity regions (LCRs) in monkeypox virus (MPXV) genome sequence 353R. Short tandem repeats (STRs) are described using nucleotide base-pair coordinates with reference to the high-quality genome (HQG) sequence (ENA Accession #OX044336). Listed are the number of repeat units, description of the sequence (with _n_ = number of repeats for this particular genome), identification of the nearest annotated orthologous poxvirus gene (OPG), type of LCR (STR or homopolymer), position of the LCR to the nearest gene, and distance of the LCR to the nearest gene. OPG notations follow the standardized nomenclature (Senkevich et al., 2021); vaccinia virus (VACV) Copenhagen strain and classical VACV gene notations are shown in addition to enable comparisons.

In general, LCRs were resolved using the assembly obtained from single-molecule sequencing and further validated using short-read sequencing since most sequences were 13– 67 bp long and therefore were covered by reads from each side or flanking region without mismatches (**Figure S1B**; **File S1)**. All LCRs (except pair 1/4 and 3) were validated this way. LCR pair 1/4 (256 bp) and LCR3 (468 bp) were only resolved with single-molecule sequencing reads due to their lengths (**Table 2**).

**Table 2.**
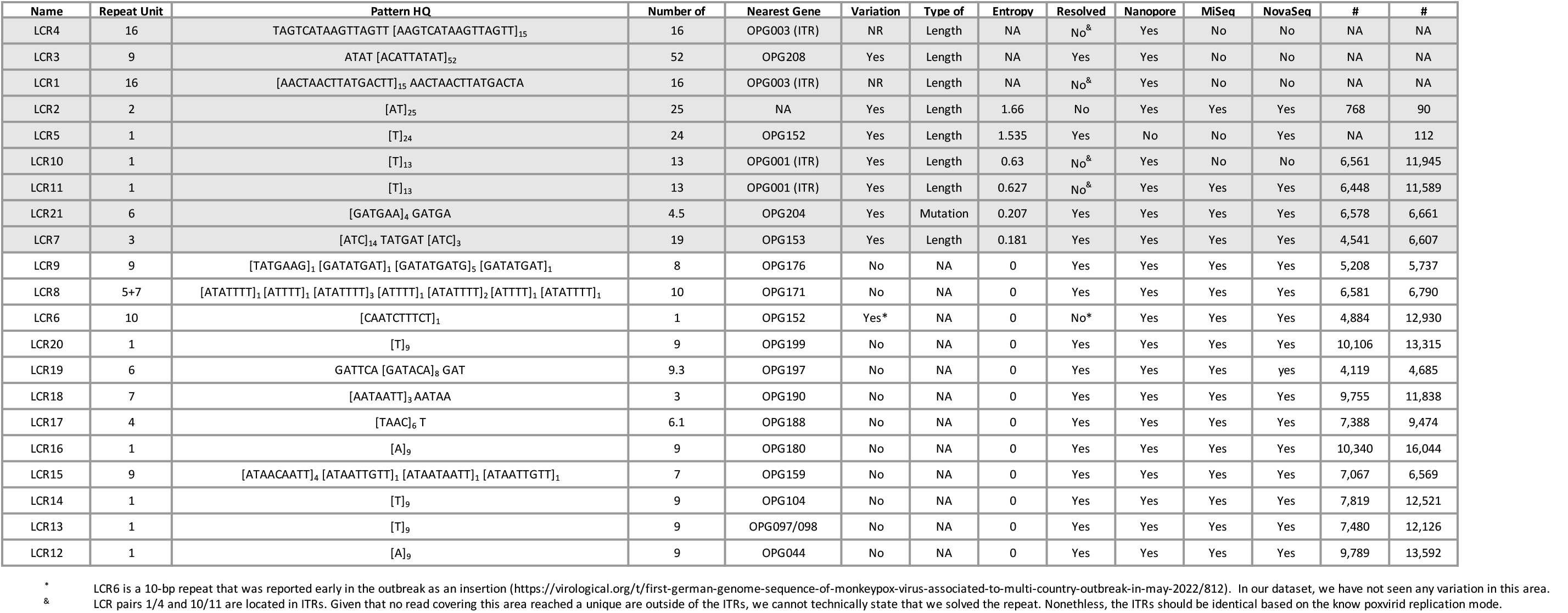
Low-complexity region (LCR) validation and entropy-level intra-host analysis in monkeypox virus (MPXV) genome sequence 353R. Listed are the type and number of supporting reads for each LCR. Definitions of quality: Yes, LCR is found entirely in the assembly in one contig; no, LCR is not assembled with the reported method. All LCRs with entropy levels above 0.15 are shaded in gray. OPG, orthologous poxvirus gene.

LCR3 contains a complex tandem repeat with the sequence ATAT [ACATTATAT]_n_. Our analysis indicated n=52. No publicly available MPXV genome sequence contains a tandem repeat of similar length. However, applying the analysis to 35 publicly available MPXV National Center for Biotechnology Information (NCBI) Sequence Read Archive (SRA) datasets of single-molecule raw reads allowed us to resolve the LCR3 of some (**Figure 2A**). Fifteen datasets revealed supporting long reads that include both LCR3 flanking regions. Interestingly, four subclade IIb lineage B.1 MPXV sequences associated with the 2022 monkeypox epidemic and available in SRA have n=54–62 repeats in LCR3. Their number of repeats separate these sequences from 2018–2019 subclade IIb lineage A sequences that have n=12–42 repeats in LCR3, indicating LCR3 as a region of genomic instability and high variability.

**Figure 2.**
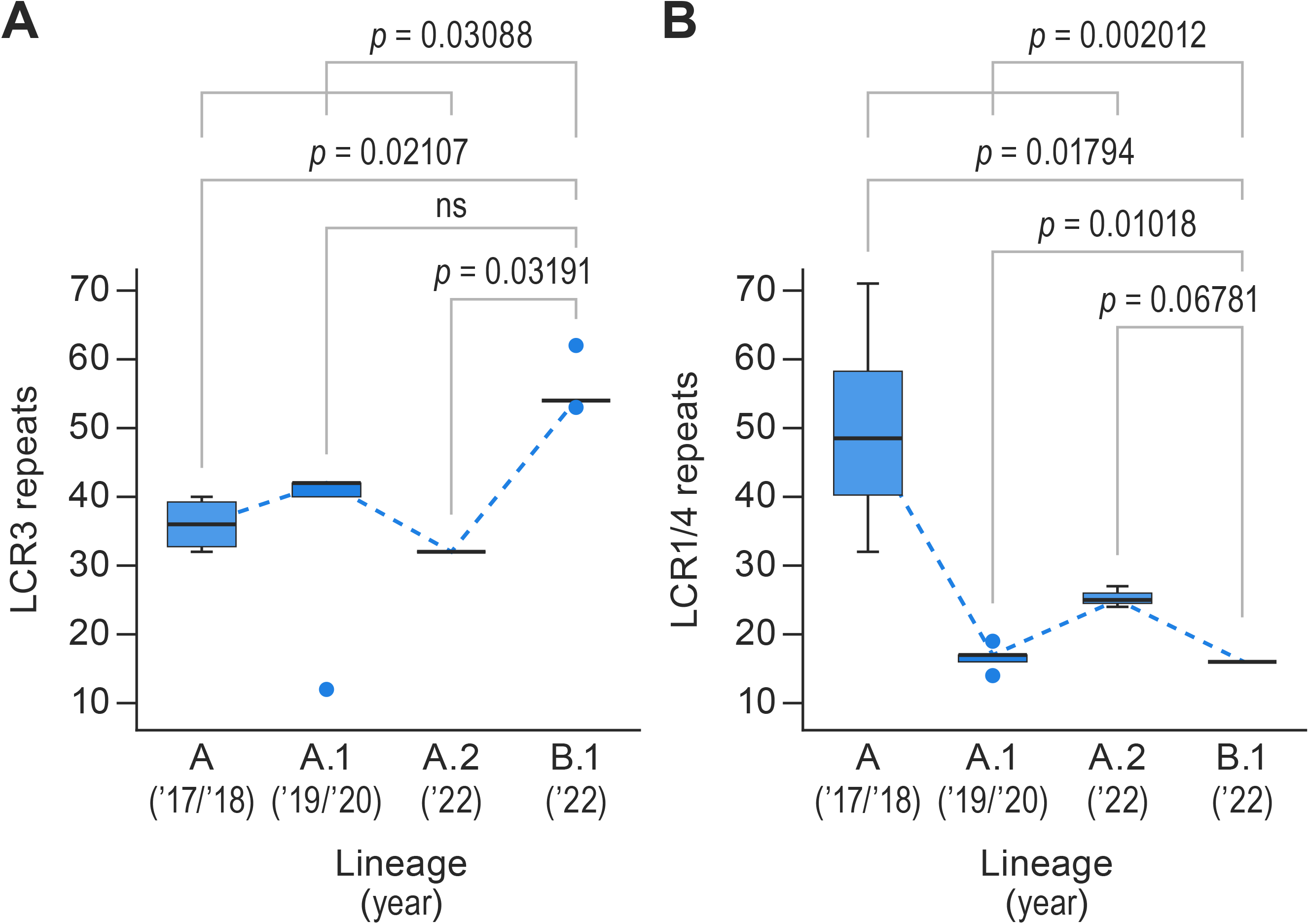
Characterization and validation of non-randomly distributed low-complexity regions (LCRs) in monkeypox virus (MPXV) genome sequence 353R. (A) LCR3 sequence validation using MPXV 353R Nanopore sequencing data and 15 additional raw data sequencing reads downloaded from the National Center for Biotechnology Information (NCBI) Sequence Read Archive (SRA). (B) LCR pair 1/4 sequence validation using MPXV 353R Nanopore sequencing data and 20 additional raw data sequencing reads downloaded from NCBI SRA. Detailed information on the represented materials, along with originator and epidemiological data, is provided in **Table S6**.

LCR pair 1/4 contains a tandem repeat with the sequence [AACTAACTTATGACTT]_n_. Our analysis indicated n=16. Instead, the sequences of the subclade IIb lineage B.1 MPXV isolate MPXV_USA_2022_MA001 and lineage A reference isolate MPXV-M5312_HM12_Rivers LCR pair 1/4 have n=8 (**Table 3**). Inclusion of the NCBI SRA datasets into the analysis confirmed the n=16 value (**Figure 2B**). In addition, the analysis revealed subclade IIb lineage-specific repeat differences in LCR pair 1/4. Lineage A.1 viruses are polymorphic, having 14 repeats (n=1 sequences), 16 (n=3), 17 (n=7), and 19 (n=1); lineage A.2 viruses have n=23–26 repeats; lineage A viruses have n=32, 43, 53, or 71 repeats. In contrast, lineage B1 viruses consistently have n=16. While LCR3 appears to have “increased” in length since the spillover, LCR pair 1/4 appears to be decreasing in length, thus behaving like an “accordion” over time.

**Table 3.**
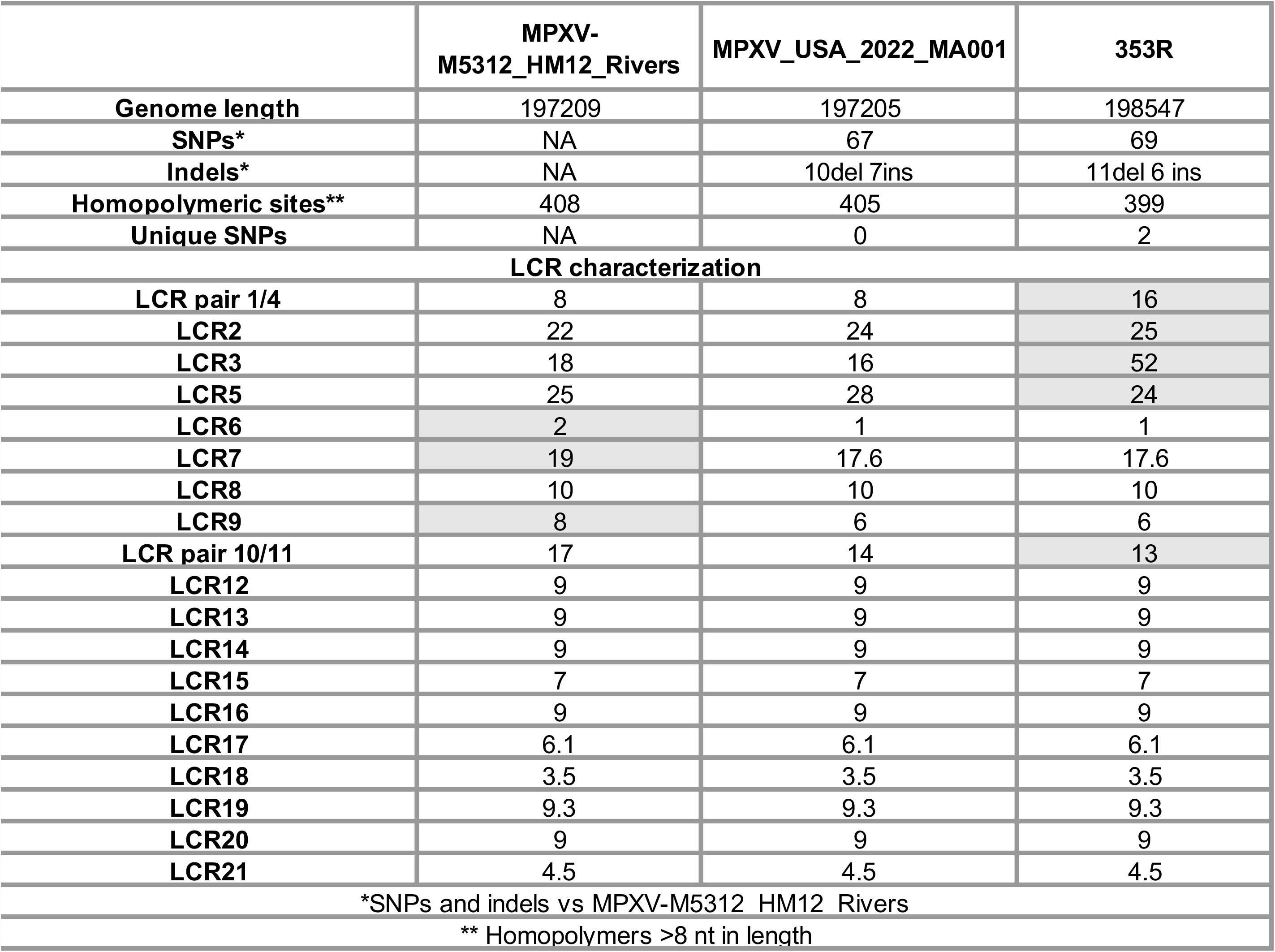
Comparison of monkeypox virus (MPXV) genome sequence 353R with reference sequences. LCR repetitions for each genome are indicated. Discrepant number (n) of LCR repeats are shaded in gray. indels, number of insertion (ins) or deletion (del) of bases; LCR, low-complexity region; SNP, single-nucleotide polymorphism.

The subclade II lineage B.1 353R and MPXV_USA_2022_MA001 genome sequences have the same 67 SNPs called against the subclade II lineage A reference isolate MPXV-M5312_HM12_Rivers genome sequence. Additionally, the 353R sequence has two additional paired SNPs in the left and right ITRs (5,595G→A; 191,615C→T compared with the MPXV-M5312_HM12_Rivers sequence) that result in the introduction of a stop codon in OPG015. We observed this variation in only two other patients among our sample set. The 353R and MPXV_USA_2022_MA001 sequences also differ by two number of insertion (ins) or deletion (del) of bases (indels) at positions 133,077 and 173,273, respectively, which correspond to differences in LCR2 and LCR5, respectively. As a result of the resolution LCRs, sequence 353R differs by 1,342 bp in genome length compared with the MPXV_USA_2022_MA001 sequence and 1,338 bp compared with the MPXV-M5312_HM12_Rivers sequence. Most of the variation is due to differences in the length of LCR pair 1/4 and LCR3, along with minor length differences in LCR2, LCR5, and LCR pair 10/11 (**Table 3**). In general, the number of repeats (n) found with the hybrid assembly approach we used here doubled the length of LCRs.

Based on the higher resolution of the MPXV 353R genome sequence, in particular regarding LCRs, we propose this sequence as the new MPXV high-quality genome (HQG) reference sequence according to the sequence quality standards defined in (Ladner et al., 2014).

### Low-complexity regions (LCRs) are non-randomly distributed in the monkeypox virus (MPXV) genome

We compared the distribution of LCRs between the different major functional protein OPG groups following the classification of (Senkevich *et al*., 2021). Differences between functional groups were statistically significant (Kruskal–Wallis test, χ^2^ *p*-value <0.001). Pairwise analysis demonstrated that the functional group “core” (orthopoxvirus genomic central conserved region) includes LCRs at a significantly lower frequency (multiple pairwise-comparison Wilcoxon test) than functional groups “ANK/PRANC” (false discovery rate [FDR]-corrected *p*-value <0.0001), “Bcl-2 domain” (corrected *p*-value = 0.04), and “accessory” (FDR-corrected *p*-value <0.0001) (**Figure 3A**). These analyses indicate that LCRs in orthopoxvirus genomes are non-randomly distributed and that there is a significant purifying selection force against introducing LCRs in central conserved region areas.

**Figure 3.**
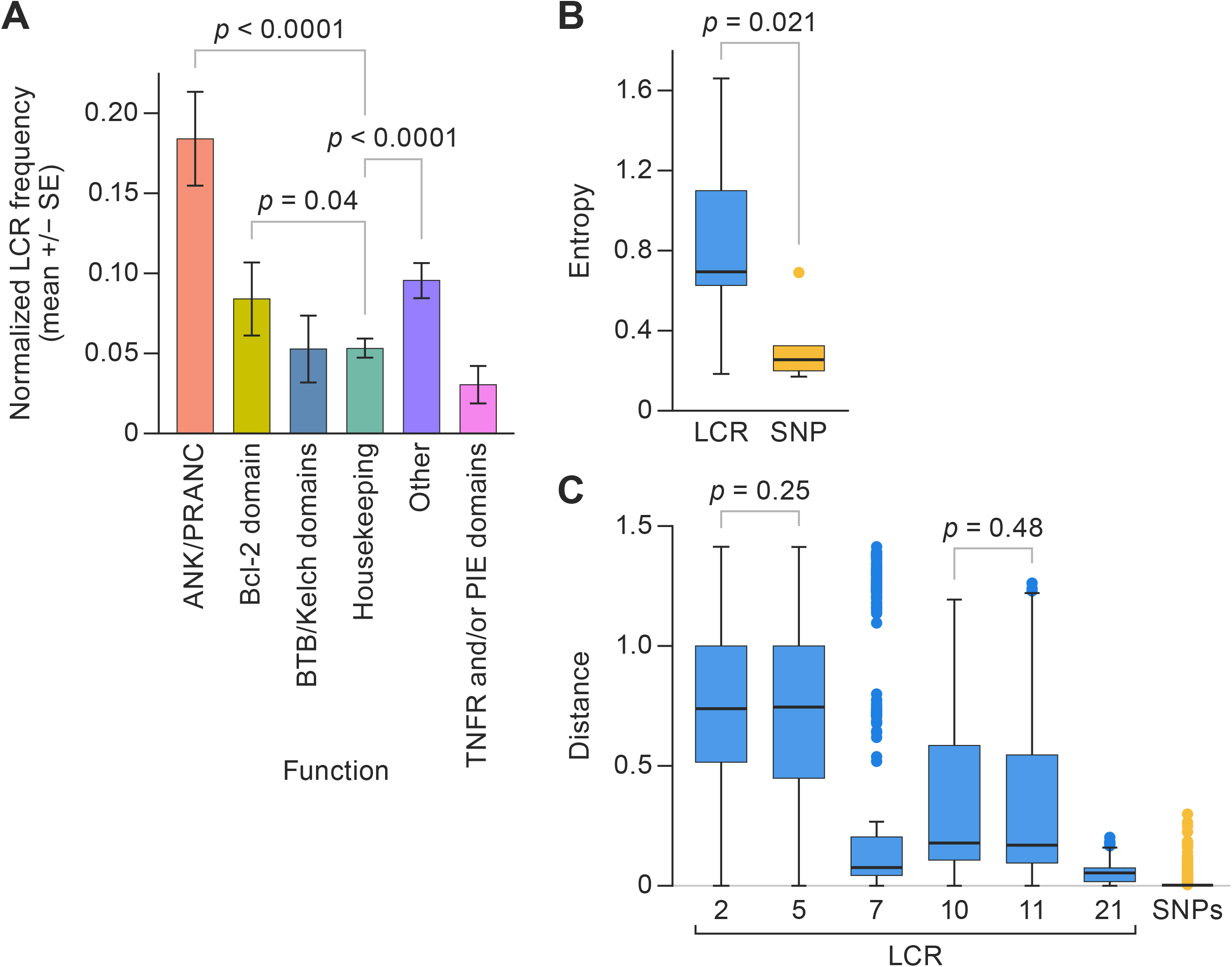
Low-complexity regions (LCRs) are non-randomly distributed in the monkeypox virus (MPXV) genome. (A) Frequencies (mean +/- standard error) at which LCRs occur in orthologous poxvirus genes (OPGs) of different functional groups. Shown are functional classes in which pairwise comparisons had a significantly different frequency than other groups (multiple pairwise Wilcoxon test, false discovery rate [FDR] corrected *p*-value <0.05). (B) Entropy value distribution for short tandem repeats (STRs) in LCRs (left) and single-nucleotide polymorphisms (SNPs; right). (C) Distributions of the pairwise inter-sample Euclidean distances for each STR in LCRs (**Table S3**). SNPs in the boxplot represent the distribution of average Euclidean distances of each variable position along the MPXV genome.

Next, we compared the degree of diversity among the 21 identified LCRs with the observed SNP variability that had been the focus of the field. In the 353R HQG sequence, LCRs 2, 5, 7, 10, 11, and 21 had intra-host genetic diversity, with entropy values that ranged from 0.18 (LCR7) to 1.66 (LCR2), with an average of 0.81 and a standard deviation (SD) of 0.64 among them (**Table 2**). Only five nucleotide positions (1,285; 6,412; 88,807; 133,894; and 145,431) had intra-host genetic diversity at the SNP level. The entropy values ranged from 0.17 (position 133,894) to 0.69 (position 6,412), with an average of 0.38 and a SD=0.21 among them. Interestingly, a student’s t-test revealed a significantly higher level of diversity in LCRs than in SNPs (*p*-value = 0.021; **Figure 3B**).

Then, we characterized, collected, and compared the allele frequencies for all LCRs from all dataset samples (**Table S3**) applying the filters described above. Our analyses revealed that the average inter-sample Euclidean distances at LCRs ranged from 0.05 (LCR21) to 0.73 (LCR2) (**Figure 3C**). We found statistically significant differences between LCRs (Kruskal–Wallis χ^2^ *p*-value <0.001). More specifically, multiple pairwise comparison Wilcoxon test results showed that all LCRs have significantly different levels of inter-sample distances (FDR-corrected *p*-values < 0.001), except in case of the LCR10 versus LCR11 (FDR-corrected *p*-value = 0.48) and LCR2 versus LCR5 (FDR-corrected *p*-value = 0.25) (**Figure 3C**). Average distances in SNPs were 0.0018–0.4168. Our randomization tests revealed that all LCRs have a significantly higher level of inter-sample diversity than the SNPs (all FDR-corrected *p*-values <0.05) (**Figure 3C**). These analyses uncovered that most of the variability in the orthopoxvirus genome is located in LCR. Consequently, we posit that studies focusing on phenotypic MPXV (and likely other orthopoxvirus) clade-/subclade-/lineage-/strain differences due to genomic sequence variation should change their focus to the study of LCR variability instead of SNP variability.

### Low-complexity regions (LCRs) might be more phylogenetically informative than single-nucleotide polymorphisms (SNPs) for inter-host sequence analysis

Analysis of only two monkeypox patient samples (353R and 349R) resulted in sufficient sequence coverage information to enable allele frequency comparison in most LCRs (**Figure 4A**). Their side-by-side comparison revealed differences in allele frequency in some of them (LCR2, LCR5, and LCR pair 10/11) (**Figure 4B**). The sequence coverage achieved with the remaining samples only enabled to unequivocally resolve an LCR subset (i.e., covering both flanking regions: LCRs 2, 5, 7, 8, 9, and pair 10/11). LCR8 and LCR9 were identical across the entire sample set. However, LCR7 and LCR pair 10/11 had considerable intra-host variation, as well as differences in the preponderant allele (LCR pair 10/11) between samples (**Figure 4C**).

**Figure 4.**
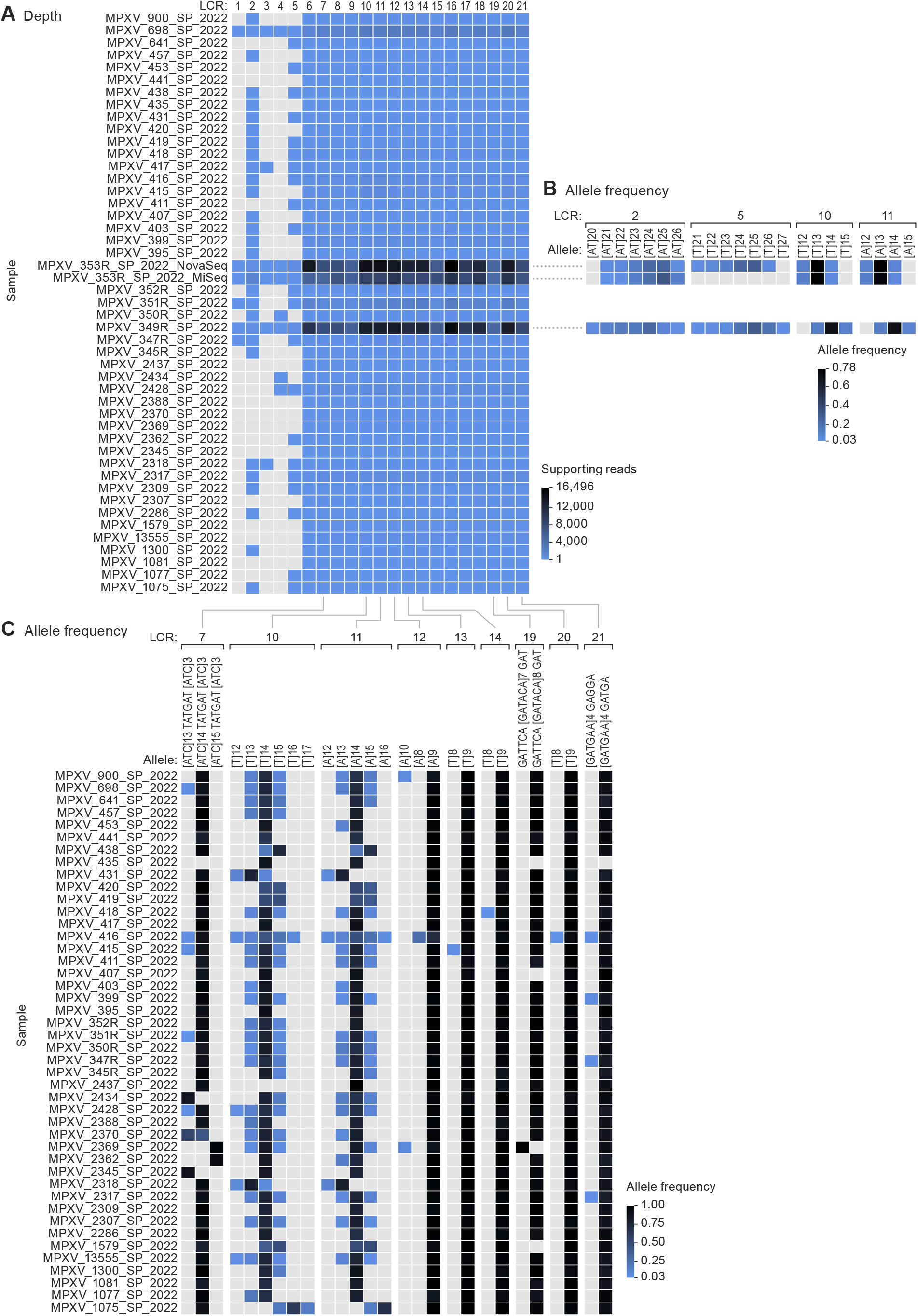
Low-complexity regions (LCRs) might be more phylogenetically informative than single-nucleotide polymorphisms (SNPs) for inter-host sequence analysis. Monkeypox virus (MPXV) population genomics within the biological specimen (intra-host) and across different specimens (inter-host). (A) Panel shading indicates the number of reads supporting each LCR for each sample. Only paired reads that include a perfect match to both flanking regions were counted; the gradient shows the maximum value in black and the minimum value (n=1) in the lightest blue. Samples without coverage are indicated in gray. (B) Comparison of LCR allele frequency for samples 353R and 349R. Only LCRs with at least 10 supporting paired reads including both flanking regions were counted; only alleles with a frequency of 0.03 or higher were considered. The gradient shows the maximum value in black and the minimum value (n=0.03) in the lightest blue. (C) Comparison of LCR allele frequency in all samples for LCRs with good coverage (7, pair 10/11, 12, 13, 14, 19, 20, and 21). Only LCRs with at least 10 supporting paired reads including both flanking regions were counted; only alleles with a frequency of 0.03 or higher were considered. The gradient shows the maximum value in black and the minimum value (n=0.03) in the lightest blue.

Phylogenetic (**Figure S2A**) and haplotype network (**Figure S2B**) analyses of the sequences determined from the monkeypox patient samples yielded limited information regarding the outbreak. Most sequences were highly similar and are, therefore, part of the basal ancestral MPXV subclade IIb lineage B1 node. Some sequences formed supported clusters: Sequences clustered into groups (**Table S4**):

- group 1 (lineage B.1): sequences from patients 395, 399, and 441 and Floridian MPXV isolate MPXV_USA_2022_FL002 (GenBank #ON676704);
- group 2 (lineage B.1): sequences from patients 347, 352R, 353R, 416, and Spanish MPXV isolate MPXV/ES0001/HUGTiP/2022 (GenBank #ON622718); all share a stop-codon mutation in OPG015;
- group 3 (lineage B.1.3): sequences from patients 22,369 formed with Slovenian MPXV isolate SLO (GenBank #ON609725.2), French isolate MPXV_FR_HCL0001_2022 (GenBank #ON622722), and 38 other sequences worldwide; defined by NBT03_gp174 mutation G190,660A, resulting in an R84K amino-acid residue change;
- group 4 (lineage B.1): sequences from patients 417 and 2,437;
- group 5 (lineage B.1.1): sequences from patients 698; 1,300; 2,388; 2,428; German MPXV isolate MPXV/Germany/2022/RKI01 (GenBank #ON637938.1) and 97 other sequences worldwide; defined by OPG094 mutation G74,360A resulting in a R194H amino-acid residue change); and
- group 6 (lineage B.1): sequences from patients 2,309 and 2,317.

Only one epidemiological link among group samples was identified; patients 395 and 399 came from sexual partners who attended events in Portugal and Spain.

In summary, at least at this time in the monkeypox epidemic, there appears to be limited value in full-genome SNP characterization for transmission analysis.

### Conservation and variation in proteins encoded by orthologous poxvirus gene (OPG) and codon usage analysis in OPG low-complexity regions (LCRs)

Analyses showed that LCRs were associated with intra-host and inter-sample variation. Although most LCRs that showed variability in our sample set (pair 1/4, 2, 3, 5, 7, pair 10/11, and 21), three (3, 7, and 21) are located in regions that, considering orthopoxvirus evolutionary history, are associated with virulence or transmission (Chen et al., 2005; Kastenmayer et al., 2014). Noteworthy, three of the 21 highly repetitive areas identified in our intra-host variation analysis (those of LCRs 5, 6, and 7) are located in a defined central conserved region of the orthopoxvirus genome between positions 130,000 and 138,000 (**Figure 1**). This region contains OPG152 (which is truncated in the MPXV genome), OPG153 (directly affected by LCR7), and OPG154. LCR7 is the only STR that is located at the center of a functional ORF). In contrast, LCR3 and LCR21 are situated in the promoter/start area, potentially modifying the ORF start site. The repeat area of LCR7 encodes a poly-D homopolymer in a nonstructured region of OPG153 (**Figure 5A**). The changes we uncovered result in the insertion of two isoleucyls. This change resembles the primary structure found in clade I viruses. In contrast, pre-2017 African subclade IIa viruses lack such insertions.

**Figure 5.**
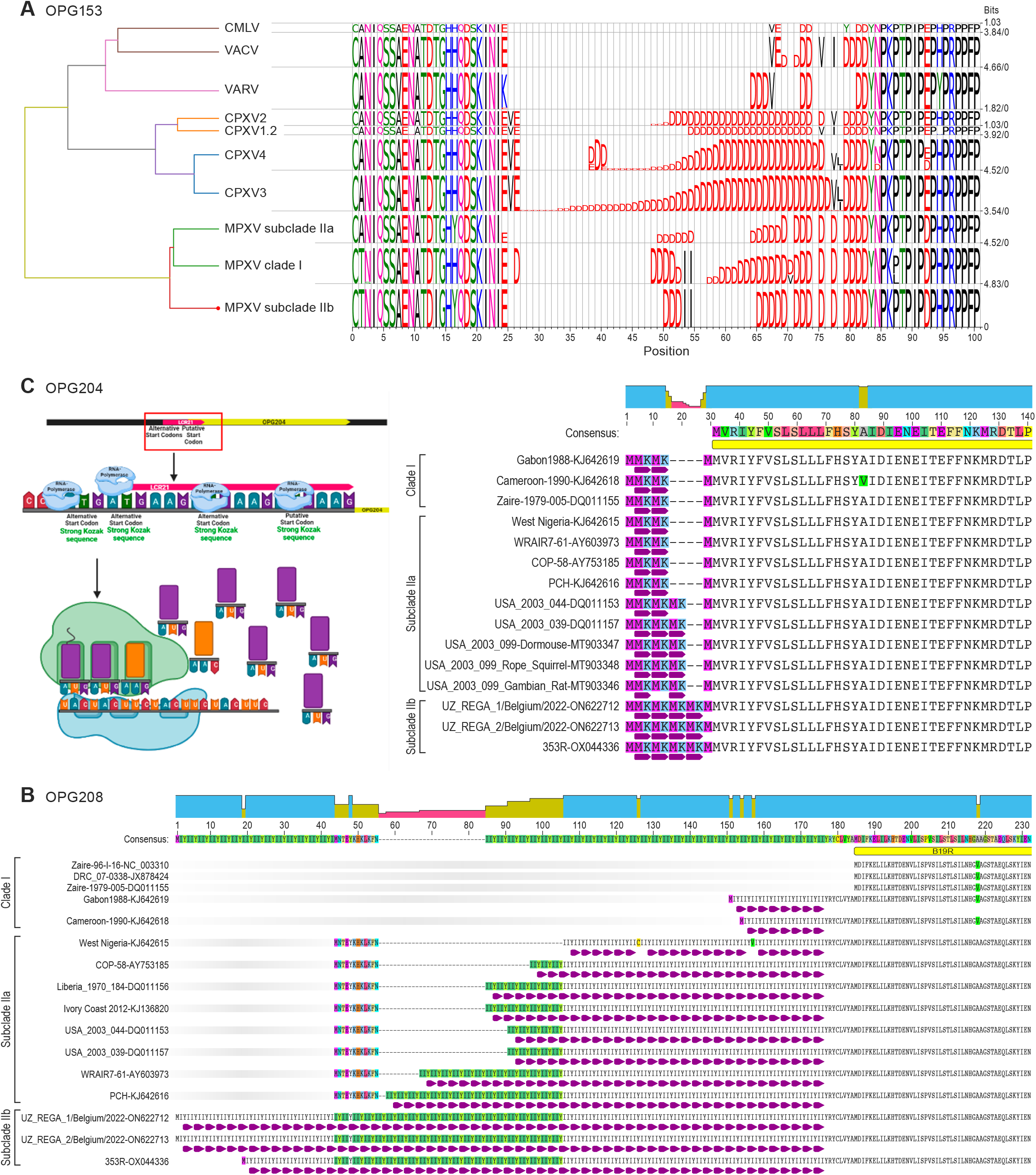
Conservation and variation in proteins encoded by orthologous poxvirus gene (OPG) and codon usage analysis in OPG low-complexity regions (LCRs) MetaLogo visualization of conserved and varying amino-acid residues in OPG-encoded proteins among monkeypox virus (MPXV) clade I, subclade IIa, and subclade IIb, with homologous and nonhomologous sites highlighted. (A) OPG153/ LCR7-derived variability. CMLV, camelpox virus; VACV, vaccinia virus; VARV, variola virus; CPXV, cowpox virus; MPXV, monkeypox virus. (B) OPG204/LCR21-derived variability; and (C) OPG208/LCR3- derived variability. Visualizations created at Biorender.com and Geneious version 2022.2 created by Biomatters.

Another region of potential functional impact is the area between 170,000 to 180,000 that includes LCRs 19, 20, 2, 21 and 3) (**Figure 1**). The LCR3 repeat [CATTATATA]_n_ is located 21 bp upstream of the putative translation start site of OPG208. Importantly, a methionyl codon is located immediately upstream of LCR3. The usage of this start codon would result in the introduction of an Ile-Ile-Tyr repeat. This codon has a medium to low probability of being used as start codon in the cognate mRNA (T base in position −3), compared to a “strong” Kozak sequence of the downstream putative start codon. Nevertheless, LCR3 remained in-frame in all clade II MPXV samples, indicating selective pressure to maintain the possibility of alternative start translation (**Figure 5B**). Interestingly, LCR3 is not located in-frame in most clade I viruses. This may be significant because OPG208 is a member of a set of genes most likely responsible for increased virulence of clade I MPXV compared to classical (pre-epidemic) clade IIa MPXV (Chen *et al*., 2005). The LCR3 tandem repeat [CATTATATA]_n_ is present with n=52, 54, and 62 copies in epidemic subclade IIb lineage B viruses (**Figure 2A**), whereas it is n=7, 37, and 27 in subclade IIa MPXV isolates Sierra Leone (GenBank #AY741551), MPXV-WRAIR7-61 (GenBank #AY603973), and MPXV-COP-58 (GenBank #AY753185) sequences, respectively (Chen *et al*., 2005), as well as n=16 for clade I MPXV isolate Zaire-96-I-16 (GenBank #AF380138). Interestingly, all publicly available subclade II lineage A single-molecule long-read sequence data imply a repeat n<40 (**Figure 2A**). Alternatively, the LCR3 repeat sequence could also alter promoter function. Interestingly, the repeat sequence also introduces codons with a low usage ratio that are not be optimized for expression in primates. The codon triplet ATA, which encodes isoleucyl, has a rare codon usage of 0.17 (Stothard, 2000) (**Figure 5B**).

Similarly, the STR downstream of LCR21 introduces a methionyl codon upstream of the putative start codon for OPG204 (**Figure 5C**). Kozak sequence analysis revealed a medium to high probability for translation compared with the putative start codon (**Figure 5C**).

The remaining LCRs (2, pair 4/1, and pair 10/11) are located downstream of known ORFs; thus, their variation is less likely to be associated with a change in phenotype.

## DISCUSSION

MPXV subclade IIb traces back to a human MPXV infection that likely occurred after spillover from a local animal reservoir in Ihie, Abia State, Nigeria, in 1971. An additional 10 human infections with MPXV of this lineage were detected through 1978, when this lineage seemed to have disappeared. However, in 2017, it reemerged in Yenagoa, Bayelsa State, Nigeria (Faye *et al*., 2018). Since then, hundreds of monkeypox cases have been reported and MPXV belonging to the IIb lineage has been sampled in several countries. But, there were no secondary cases prior to the 2022 epidemic (Cohen-Gihon et al., 2020; Ng et al., 2019; Vaughan et al., 2018; Yong et al., 2020).

Subclade IIb viruses cause monkeypox that presents differently than the classical disease caused by clade I and subclade IIa viruses, i.e., subclade IIb infections are associated with higher prevalence among adults rather than adolescents, are predominant in the MSM community, and are efficiently transmitted human-to-human from localized infectious skin lesions rather than requiring disseminated infection (Bunge *et al*., 2022; Otu *et al*., 2022; Thornhill *et al*., 2022; Ulaeto *et al*., 2022; Vusirikala *et al*., 2022).

Comparative genomics demonstrated obvious relationships between orthopoxvirus genotype and phenotype, driven by selective pressure from hosts (Baroudy and Moss, 1982; Chen *et al*., 2005; Esposito et al., 2006; Gubser et al., 2004; Hendrickson et al., 2010; Kugelman *et al*., 2014; Shchelkunov, 2012; Shchelkunov et al., 2001). Consequently, it was expected that increased MPXV genotype IIb human-to-human transmission would go hand in hand with genotypic changes. However, since orthopoxviruses genomes are organized in redundant ways (Bratke and McLysaght, 2008; Elde *et al*., 2012; McLysaght et al., 2003; Senkevich *et al*., 2020), genotypic changes were expected to be modulating, rather than radical.

Thus far, genomic characterization of the 2022 epidemic has focused on describing its evolutionary history and tracking MPXV introductions into western countries. The 2022 MPXV cluster diverges from its predecessor viruses by an average of 50 SNPs. Of these, the majority (n=24) are non-synonymous mutations with a second minority subset of synonymous mutations (n=18) and a few intergenic differences (n=4) (Isidro et al., 2022). A strong mutational bias was mainly attributed to the potential action of apolipoprotein B mRNA-editing catalytic polypeptide-like 3 (APOBEC3) enzymes (O’Tool and Rambaut, 2022). Genetic variation, including deletion of immunomodulatory genes, also occurred (Jones et al., 2022). MPXV sublineages mostly represent very small variations, usually characterized by one or two SNP differences to basal nodes in phylogenetic analyses (Nextstrain, 2022). Four of the MPXV genome sequences determined in this study can be assigned to global lineage B.1.1, one sequence could be assigned to global lineage B.1.3, and the remaining new sequences belong to lineage B1. We detected additional clusters that were also defined by few SNPs, but only in one case could we identify an epidemiological link. Thus, there appears to be a limited relationship between SNPs and epidemiology, which might hint to sequencing errors. Consequently, MPXV genomic epidemiology might need a change of focus.

Our analyses located considerable MPXV genomic variability in areas previously considered of poor informative value, i.e., in LCRs. Because LCR entropy is significantly higher than that of SNPs and LCRs are not randomly distributed in defined coding areas in the genome, person-to-person transmission-associated changes were observed in the immunomodulatory region (Kugelman *et al*., 2014), and genomic accordions are a rapid path for adaptation of orthopoxvirus during serial passaging (Elde *et al*., 2012; Senkevich *et al*., 2020), we posit that LCR changes might be associated with MPXV transmissibility differences over time.

Eight LCRs had evident signs of intra-host and inter-sample variation (pair 1/4, 2, 3, 5, 6, 7, pair 10/11, 21). Five of them (5, 6, 7, 3, and 21) were co-located in two areas of the MPXV genome: base pairs 130,000–135,000 (5, 6, and 7) which are in the central conserved region of the OPXV genome in which most “housekeeping” genes are located; and base pairs 170,000–180,000 (3 and 21), which are located in the immunomodulatory area (**Figure 1**). Three of those LCRs are located inside the putative translated regions of genes OPG153, OPG204, and OPG208. Changes in OPG204 and OPG208 are located near the N-terminal region and might involve modulating the expression of translation. The changes observed in OPG153 stand out as they are located inside a region that is under high selective pressure for transmission in a “housekeeping” orthologous poxvirus gene, which is involved in virion attachment and egress (Senkevich *et al*., 2021). Thus, we urge that these areas be scrutinized for changes that might affect the MPXV interactome.

The OPG153 repeat results in a poly-Asp amino-acid homopolymer string (**Figure 5A**); the LCR repeat in OPG208 results in an Ile-Ile-Tyr repeat (**Figure 5B**); and the N-terminal domain variation in OPG204 results in a Met-Lys repeat (**Figure 5C**). Self-association guided by stretches of single amino-acid residue repeats may lead to the formation of aggregates (Oma et al., 2004). Many human diseases are associated with detrimental effects of homopolymers (Gatchel and Zoghbi, 2005; Shoubridge et al., 2007). Expansion or contraction of the repeat may increase self-attraction and trigger disease. These homopolymers also regulate the activity of transcription factors (Gemayel et al., 2015) or direct proteins to different cellular compartments (Oma *et al*., 2004; Salichs et al., 2009). Modulation of ORF translation via MPXV LCR-like repeats also has been described for various microbes. For instance, the functionality of baker’s yeast proteins Flo1p and Flo11p is proportionally modulated by the repeat length of their N-terminal regions. A relatively small change in the number of tandem repeats is crucial for yeast adaptation to a new environment (Fidalgo et al., 2006; Verstrepen et al., 2005). Another example is glutamic acid-rich protein (GARP) of plasmodia, which contains repetitive sequences that direct the protein to the periphery of the infected erythrocyte. At least nine other exported plasmodium proteins target the periphery of the erythrocyte using this strategy. Interestingly, the lengths of the tandem repeats vary among plasmodium strains (Davies et al., 2017; Davies et al., 2016).

Protein translation rates are in part regulated by the availability of mRNA codon-cognate aminoacylated tRNAs. Homopolymers composed of codons for rare tRNAs directly diminish translation via tRNA depletion. The non-optimal tyrosyl codons in MPXV LCRs 3 and 21 suggest such translation modulation for OPG208 and OPG204, respectively. The protein encoded by OPG208, B19R (Cop-K2L, SPI-1) is a serine protease inhibitor-like protein that functions as an inhibitor of apoptosis in VACV-infected cells (Kettle et al., 1995; Kotwal and Moss, 1989), which could prevent VACV proliferation and protect nearby cells (Brooks et al., 1995; Jorgensen et al., 2017). Consequently, MPXV B19R is now considered a potential MPXV virulence marker (Chen *et al*., 2005). The protein encoded by OPG204, B16R, is a secreted decoy receptor for interferon type I (Colamonici et al., 1995; Hernáez et al., 2018). We did not observe any repeat number changes OPG204-associated LCR21, but SNPs in clade I, subclade IIa, and subclade IIb result in alternative translational start sites followed by a, suggesting these SNPs could have direct effects on OPG204 translation.

The most intriguing finding from our dataset involves LCR7 in OPG153. The OPG153 expression product (A26L) attaches orthopoxvirus particles to laminin (Chiu et al., 2007) and regulates orthopoxvirus particle egress (Howard et al., 2008; Kastenmayer *et al*., 2014; Liu et al., 2014; Ulaeto et al., 1996), thereby modulating key steps in the virus lifecycle. OPG153 is unique as it is the central conserved region gene that has been “lost” the most times during orthopoxvirus evolution (Senkevich *et al*., 2021). Inactivation of OPG153 genes by frameshift mutations occurs rapidly in experimental orthopoxvirus evolution models (Senkevich *et al*., 2020), resulting increased virus replication levels, changes in particle morphogenesis, decreased particle-to-PFU ratios, and differences in pathogenesis (Kastenmayer *et al*., 2014; Senkevich *et al*., 2020). Finally, A26L is the main target of the host antibody response to orthopoxvirus infection (Keasey et al., 2010; Pugh et al., 2016). Thus, any genomic change that modulates OPG153 is likely of significance. LCR7 encodes a poly-D non-structured region that is conserved among orthopoxviruses A26Ls; however, its length is highly variable. In mammals, poly-D stretches appear to provide functionality to asporin, a small leucin rich repeat proteoglycan that also possesses a unique stretch of aspartyls at its N terminus (Henry et al., 2001), associated with calcium-binding (Zhu et al., 2018). Interestingly, orthopoxviruses that form a dense protein matrix within the cytoplasm called A-type inclusions (ATIs), such as MPXV, generically have very long poly-D stretches in this region, whereas orthopoxviruses that do not form ATIs, such as VACV and VARV, have reduced their LCR7 poly-D stretches to four Ds. Even among MPXV clades, patterns are observable. Subclade IIa viruses have an extended 21 amino-acid residue poly-D stretch, whereas clade I and subclade IIb viruses have poly-D stretches with two inserted isoleucyls. Intriguingly, both insertions result from the incorporation of the same “ATCATA” nucleotide insertion in the “GAT” repetitive stretch.

In summary, our findings expand the concept of genome accordions as a simple and recurrent mechanism of adaptation on a genomic scale in orthopoxvirus evolution. A consequence of this broadening is the recognition that MPXV genome LCRs might hold the key to improved understanding of current monkeypox epidemiology and clinical presentation. A new standardized approach to generate and analyze sequencing data via prioritizing LCR characterization and subsequent functional mechanism-of-action studies are warranted.

## MATERIALS AND METHODS

### Study population

This study includes confirmed human monkeypox cases diagnosed from May 18 to July 14, 2022, at the Centro Nacional de Microbiología (CNM), Instituto de Salud Carlos III, Madrid, Spain. The study was performed as part of the public health response to the current monkeypox epidemic by the Spanish Ministry of Health. Sample information is listed in **Tables S1** and **S5**.

The samples used in this work were obtained in the context of the Microbiological Surveillance and Diagnosis Program for the Monkeypox Outbreak of the Centro Nacional de Microbiología, Instituto de Salud Carlos III. The study was based on routine testing, did not involve any additional sampling or tests and stored RNA extracts were used, so specific ethical approval was not required for this study. All sequenced viruses corresponded to those to patients that gave consent to be analyzed for diagnosis or surveillance purposes.

### Study sample processing

Swabs of vesicular lesions from study patients in viral transport media were sent refrigerated to CNM. Nucleic acids were extracted at CNM using either QIAamp MinElute Virus Spin (DNA) or QIAamp Viral RNA Mini kits (Qiagen, Germantown, MD, USA) according to the manufacturer’s recommendations. Inactivation of samples was conducted in a certified class II biological safety cabinet in a biosafety level (BSL) 2 laboratory using BSL-3 best practices with appropriate personal protective equipment.

### Monkeypox virus (MPXV) laboratory confirmation

MPXV detection by PCR in a sample was considered laboratory confirmation and resulted in inclusion of the swab in the study. A previously described orthopoxvirus-generic real-time PCR (qPCR) was used for screening (Fedele et al., 2006). A previously described conventional validated nested PCR targeting OPG002 (encoding a TFN receptor) was used for results confirmation (Sánchez-Seco et al., 2006).

### MPXV genome sequencing

Sequencing libraries were prepared with a tagmentation-based Illumina DNA Prep kit (Illumina, San Diego, CA, USA) and run in a NovaSeq 6000 SP Reagent Kit (Illumina) flow cell using 2×150 paired-end sequencing. To improve assembly quality, the library from swab 353R, an unpassaged vesicular fluid from a confirmed case, was also run in a MiSeq Reagent Kit v3 (Illumina) flow cell using 2 x 300 paired-end sequencing. Additionally, sample 353R was also analyzed by single-molecule methods using Nanopore sequencing (Oxford Nanopore Technologies, Oxford, UK). For Nanopore sequencing, 210 ng of DNA was extracted from swab 353R and used to prepare a sequence library with a Rapid Sequencing Kit (Oxford Nanopore Technologies); the library was analyzed in an FLO-MIN106D (Oxford Nanopore Technologies) flow cell for 25 h. The process rendered 1.12 Gb of filter-passed bases.

### Bioinformatics

#### *De novo* assembly and annotation of subclade II lineage B.1 MPXV genome sequence 353R

Due to the high yield of MPXV genomic material in a preparatory run, swab 353R was selected as source material for the determination of an MPXV high-quality genome (HQG) sequence. Single-molecule long-sequencing reads were preprocessed using Porechop v0.3.2pre (Wick et al., 2017) with default parameters. Reads were *de novo* assembled using Flye v2.9-b1768 (Kolmogorov et al., 2019) in single-molecule sequencing raw read mode with default parameters, resulting in one MPXV contig of 198,254 bp. Short 2×150 sequencing reads were mapped with Bowtie2 v2.4.4 (Langmead and Salzberg, 2012) against the selected contig, and resulting BAM files were used to correct the assembly using Pilon v1.24 (Walker et al., 2014). At this intermediate step, this corrected sequence was used as a reference in the nf-core/viralrecon v2.4.1 pipeline (Patel et al., 2022) for mapping and consensus generation with short sequencing reads. The allele frequency threshold of 0.5 was used for including variant positions in the corrected contig.

Short MiSeq 2×300 and NovaSeq 2×150 sequencing reads were also assembled *de novo* using the nf-core/viralrecon v2.4.1 pipeline, written in Nextflow (Di Tommaso et al., 2017) in collaboration between the nf-core community (Ewels et al., 2020) and the Unidad de Bioinformática, Instituto de Salud Carlos III, Madrid, Spain (https://github.com/BU-ISCIII). FASTQ files containing raw reads were quality controlled using FASTQC v0.11.9 (Andrews, 2010). Raw reads were trimmed using fastp v0.23.2 (Chen et al., 2018). The sliding-window quality-filtering approach was performed, scanning the read with a 4-base-wide sliding window and cutting 3**’** and 5**’** base ends when average quality per base dropped below a Qphred33 of 20. Reads shorter than 50 bp and reads with more than 10% read quality under Qphred 20 were removed. Host genome reads were removed via kmer-based mapping of the trimmed reads against the human genome reference sequence GRCh38 (https://www.ncbi.nlm.nih.gov/data-hub/genome/GCF_000001405.26/) using Kraken 2 v2.1.2 (Wood et al., 2019). The remaining non-host reads were assembled using SPADES v3.15.3 (Antipov et al., 2016; Prjibelski et al., 2020) in rnaviral mode. A fully ordered MPXV genome sequence was generated using ABACAS v1.3.1 (Assefa et al., 2009), based on the MPXV isolate MPXV_USA_2022_MA001 (Nextstrain subclade IIb lineage B.1) sequence (GenBank #ON563414.3) (Gigante et al., 2022). The independently obtained *de novo* assemblies and reference-based consensus genomes obtained from swab 353R were aligned using MAFFT v7.475 (Katoh et al., 2019) and visually inspected for variation using Jalview v2.11.0 (Waterhouse et al., 2009).

#### Systematic identification of low-complexity regions in orthopoxvirus genomes

Detection of short tandem repeats (STRs) in the HQG sequence and other orthopoxvirus genomes was performed with Tandem repeats finder (Benson, 1999), using default parameters. Briefly, the algorithm works without the need to specify either the pattern or its length. Tandem repeats are identified considering percent identity and frequency of insertion (ins) or deletion (del) of bases (indels) between adjacent pattern copies, using statistically based recognition criteria. Since Tandem repeats finder does not detect single-nucleotide repeats, we developed an R script to systematically identify homopolymers of at least 9 nucleotide residues in all available orthopoxvirus genome sequences. STRs and homopolymers were annotated as low-complexity regions (LCRs).

#### Curation of low-complexity regions in the MPXV high-quality virus genome sequence

We curated LCRs in the HQG sequence using a modified version of STRsearch (Wang et al., 2020). Once provided with identifying flanking regions, STRsearch performed a profile analysis of STRs in massively parallel sequencing data. To ensure high-quality characterization of LCR alleles, we modified the script (https://github.com/BU-ISCIII/MPXstreveal) to complement reverse reads that map against the reverse genome strand according to their BAM flags. In addition, output was modified to add information later accessed by a custom Python script to select only reads containing both LCR flanking regions. All LCRs in the HQG sequence were manually validated using STRsearch results and *de novo* assemblies obtained from all sequencing approaches. When an LCR was only resolved by single-molecule long-sequencing technologies (LCR pair 1/4 and LCR3), we also analyzed publicly available data by downloading all single-molecule long-sequencing data from the National Center for Biotechnology Information (NCBI) Sequence Read Archive (SRA) (https://www.ncbi.nlm.nih.gov/sra) as of August 10, 2022, and analyzed the data according to **File S1**.

#### Final MPXV high-quality virus genome sequence assembly

The consensus genome constructed with the nf-core/viralrecon v2.4.1 pipeline using the corrected *de novo* contig as stated above, along with the resulting curated and validated consensus LCRs, were used to build the final HQG reference sequence using a custom Python script. The resulting HQG is available from the European Nucleotide Archive (#OX044336.2).

#### Generation of MPXV high-quality virus genome reference-based consensus sequence for all other samples

For the remaining specimens, sequencing reads were analyzed for MPXV genome sequence determination using the nf-core/viralrecon v2.4.1 pipeline. Trimmed reads were mapped with Bowtie2 v2.4.4 against the HQG sequence and the sequence of subclade II lineage A MPXV isolate M5312_HM12_Rivers (GenBank #MT903340.1) (Mauldin et al., 2022). Picard v2.26.10 (The Broad Institute, 2018) and SAMtools v1.14 (Li et al., 2009) were used to generate MPXV genome mapping statistics. iVar v1.3.1 (Grubaugh et al., 2019), which calls for low-frequency and high-frequency variants, was used for variant calling. Variants with an allele frequency higher than 75% were kept to be included in the consensus genome sequence. BCFtools v1.14 (Danecek et al., 2021) was used to obtain the MPXV genome sequence consensus with filtered variants and masked genomic regions with coverage values lower than 10X. All variants, included or not, in the consensus genome sequence, were annotated using SnpEff v5.0e (Cingolani et al., 2012b), and SnpSift v4.3 (Cingolani et al., 2012a). Final summary reports were created using MultiQC v.1.11 (Ewels et al., 2016). Consensus genome sequences were analyzed with Nextclade v2.4.1 (Aksamentov et al., 2021) using the “MPXV (All clades)” dataset (timestamp 2022-08-19T12:00:00Z). Raw reads and consensus genomes are available from the European Nucleotide Archive (#ERS12168855–ERS12168865, #ERS12168867, #ERS12168868, #ERS13490510–ERS13490543).

#### Intra-host and inter-host allele frequency analyses

Intra-host genetic entropy (defined as - sum(Xi*log(Xi)), in which Xi denotes each of the allele frequencies in a position) was calculated according to the single-nucleotide polymorphisms (SNP) frequencies of each position along the genome using nf-core/viralrecon v2.4.1 pipeline results. Similarly, genetic entropy for each LCR was calculated considering the frequencies of repeat lengths.

LCR intra-host and inter-host variations in the sample set were analyzed using the modified version of STRsearch. As a filter for quality for this analysis, STRsearch results (**Table S5**) were filtered, keeping alleles with at least 10 reads spanning the region and allele frequency above 0.03. Quality control and allele frequency graphs were created using a customized R script.

Pairwise genetic distances between samples were calculated as Euclidean distances (defined as /X-Y/=sqrt(sum(xi-yi)^2), in which xi and yi are the allele frequencies of sample X and Y at a given position, respectively), thus accounting for the major and minor alleles at each analyzed position. Distances were calculated individually for each variable LCR (STRs 2, 5, 7, 10, 11, and 21) and for each of all 5,422 SNPs showing inter-sample variability (compared to MPXV-M5312_HM12_Rivers). The distributions of inter-sample distances were compared between LCRs using a Kruskal–Wallis test (χ^2^ *p*-values) followed by multiple pairwise-comparison between groups (Wilcoxon test), with *p*-values subjected to the false discovery rate (FDR) correction. A randomization test was used to test whether inter-sample variability in LCRs is higher than that in SNPs: first, the average Euclidean distance for each LCR and each SNP position was calculated; then, the average value of each LCR was compared to a random sample of 1,000 values from the distribution of mean distances from the SNPs along the genome. The *p*-value was calculated from the percentage of times that the mean of the LCR was higher than the randomly taken values from the SNPs.

#### Phylogenetic analysis of the MPXV central conserved region

Variant calling and SNP matrix generation was performed using Snippy v4.4.5 (Seeman, 2015), including sequence samples and representative MPXV genome sequences downloaded from GenBank (**Table S5**). The SNP matrix with both invariant and variant sites was used for phylogenetic analysis using IQ-Tree 2 v. 2.1.4-beta (Minh et al., 2020) via predicted model K3Pu+F+I and 1,000 bootstrap replicates. A phylogenetic tree was visualized and annotated using iTOL v6.5.8 (Letunic and Bork, 2021). The SNP matrix was also used for generating the haplotype network using PopArt v1.7 (Bandelt et al., 1999).

#### Selected MPXV ORF analysis

Representative orthopoxvirus genomes (Senkevich *et al*., 2021) were downloaded from GenBank together with the consensus genome sequences from the specimens analyzed in this study (**Table S5**). MPXV genomes were assigned to clades and lineages according to the most recent nomenclature recommendations according to Nextstrain (Nextstrain, 2022) using Nextclade v2.4.1. Annotations from RefSeq #NC_063383.1 (subclade II lineage A MPXV virus isolate MPXV-M5312_HM12_Rivers) GFF file were transferred to all FASTA genome sequences using Liftoff v1.6.3 (Shumate and Salzberg, 2021). OPG153 was extracted using AGAT v0.9.1 (Dainat et al., 2022) and multi-FASTA files were generated for each group and gene. OPG204 and OPG208 alternative annotation start site ORFs were re-annotated in Geneious Prime (Biomatters, San Diego, CA, USA), and extracted as new alignments. We used MUSCLE v3.8.1551 for aligning each multi-FASTA file and Jalview v2.11.0 for inspecting and editing the alignments. Finally, MetaLogo v1.1.2 (Chen et al., 2022) was used for creating and aligning the sequence logos for each orthopoxvirus group of the OPG153/LCR7, OPG204/LCR21, and OPG208/LCR3 areas.

#### Comparison of LCR frequencies in protein functional groups

The potential biological impact of LCRs was evaluated by mapping the frequency and location of STRs and homopolymers in the orthopoxvirus genome and considering the biological function of the affected genes. The frequency of inclusion of LCRs between distinct functional groups of genes was compared as previously described (Senkevich *et al*., 2021). Orthopoxviruses (n=231, Akhmeta virus [AKMV]: n=6 sequences; alaskapox virus [AKPV]: n=1; cowpox virus [CPXV]: n=82; ectromelia virus [ECTV]: n=5; MPXV: n=62; VACV: n=18; VARV: n=57) include 216 functionally annotated OPGs classified in 5 categories (“Housekeeping genes/Core” ANK/PRANC family, Bcl-2 domain family, PIE family, and “Accessory/Other” [e.g., virus-host interacting genes]). The frequency was calculated after normalizing count numbers with the sample size of the OPG alignment. Statistical analysis of the significance of differences was performed by applying a Kruskal–Wallis test (χ^2^ *p*-values) followed by a non-parametric multiple pairwise comparison between groups (Wilcoxon test), with *p*-values subjected to FDR correction.

## ACKNOWLEDGMENTS

We would like to thank the work of the Rapid Response Unit of the National Center for Microbiology, especially MªJosé Buitrago, and Cristobal Belda, ISCIII General Director. We also thank Anya Crane (Integrated Research Facility at Fort Detrick, National Institute of Allergy and Infectious Diseases, National Institutes of Health) for critically editing the manuscript and Jiro Wada (Integrated Research Facility at Fort Detrick, National Institute of Allergy and Infectious Diseases, National Institutes of Health) for helping with figure preparation.

The work for this study at Instituto de Salud Carlos III was partially funded by Acción Estratégica “Impacto clínico y microbiológico del brote por el virus de la viruela del mono en pacientes en España (2022): proyecto multicéntrico MONKPOX-ESP22” (CIBERINFEC). The work for this study at the Department of Microbiology, Icahn School of Medicine at Mount Sinai as part of Global Health Emerging Pathogen Institute activities was funded by institutional funds (G.P.) from the Department of Microbiology, Icahn School of Medicine at Mount Sinai in support of Global Health Emerging Pathogen Institute activities.

This work was also supported in part through Laulima Government Solutions, LLC, prime contract with the U.S. National Institute of Allergy and Infectious Diseases (NIAID) under Contract No. HHSN272201800013C. J.H.K. performed this work as an employee of Tunnell Government Services (TGS), a subcontractor of Laulima Government Solutions, LLC, under Contract No. HHSN272201800013C.

Opinions, interpretations, conclusions, and recommendations are those of the authors and are not necessarily endorsed by the U.S. Army.

The views and conclusions contained in this document are those of the authors and should not be interpreted as necessarily representing the official policies, either expressed or implied, of the U.S. Department of Health and Human Services or of the institutions and companies affiliated with the authors, nor does mention of trade names, commercial products, or organizations imply endorsement by the U.S. Government.

## AUTHOR CONTRIBUTIONS

Conceptualization, S. M., S.V., A. N., I. J., M.S.L., A.G.S., I.C., M.S.S., G.P.

Methodology, S.M., S.V., A.N., J.A.P.G., S.V.F., A.Z., J. H. K., M.S.L., N.D., J.R.K., E.G., S.G., G.P.

Investigation, S.M., S.V., A.N., J.A.P.G., S.V.F., A.Z., E.O., O.A., A.M.G., A.D.I., V.E., C.G., F.M., P.S., M.T., A.V., J.C.G., I.T., M.C.R., L.M., M.L., A.G., L.C., A.G., J.C., L.H., P.J.S., M.L.N.R., I.J., M.E.A.A., C.L., L.R., I.E., M.S., M.A.M., J.H.K., M.S.L., N.D.P., J.R.K., E.G., S.G., A.G.S., I.C., M.S.S., G.P.

Formal analysis, S.M., S.V., A.N., J.A.P.G., S.V.F., J.H.K., M.S.L., N.D.P., J.R.K., E.G., I.C., M.S.S, G.P.

Writing – original draft, S.M., S.V., A.N., G.P.

Writing – review & editing, S. M., S.V., A. N., A.G.S., I.C., M.S.S., G.P.

Visualization, S.M., S.V., A.N., G.P.

Supervision, A.G.S., I.C., M.S.S., G.P.

Resources, A.G.S., I.C., M.S.S., G.P.

Funding Acquisition, A.G.S., I.C., M.S.S., G.P.

## DECLARATION OF INTERESTS

A.G.-S. has consulting agreements for the following companies involving cash and/or stock: Castlevax, Amovir, Vivaldi Biosciences, Contrafect, 7Hills Pharma, Avimex, Vaxalto, Pagoda, Accurius, Esperovax, Farmak, Applied Biological Laboratories, Pharmamar, Paratus, CureLab Oncology, CureLab Veterinary, Synairgen, and Pfizer, outside of the reported work. A.G.-S. has been an invited speaker in meeting events organized by Seqirus, Janssen, Abbott, and Astrazeneca. A.G.-S. is inventor on patents and patent applications on the use of antivirals and vaccines for the treatment and prevention of virus infections and cancer, owned by the Icahn School of Medicine at Mount Sinai, New York, outside of the reported work.

The authors declare no competing interests.

## RESOURCE AVAILABILITY

### Lead Contact

Further information and requests for resources and reagents should be directed to and will be fulfilled by the lead contact, Gustavo Palacios.

### Materials Availability

This study did not generate new unique reagents.

### Data and Code Availability

All scripts and codes used for this study can be found at github repository: (https://github.com/BU-ISCIII/MPXstreveal).

## SUPPLEMENTAL FIGURE LEGENDS

**Figure S1. Examples for read mapping artefacts and correction in monkeypox virus (MPXV) genome low-complexity regions (LCRs)**

(A) LCR2 alignment highlighting differences compared with various consensus sequences. (B) LCR7 alignment highlighting differences of results obtained using three sequencing platforms compared to the subclade II lineage A monkeypox reference isolate MPXV-M5312_HM12_Rivers sequence.

**Figure S2. Phylogenetic analysis of monkeypox virus (MPXV)**

(A) Phylogenetic maximum-likelihood (ML) tree showing monkeypox virus (MPXV) subclade IIb single-nucleotide polymorphism (SNP) clustering. Bootstrap supports >60 are indicated by labels with their number of supports. (B) Haplotype network showing SNP differences among samples included in the phylogenetic tree. Details on groups can be found in **Table S4**.

**Figure S3. Conservation and variation in proteins encoded by orthologous poxvirus gene (OPG) 208.** (A) MetaLogo visualization of conserved and varying amino-acid residues in OPG-encoded proteins among monkeypox virus (MPXV) clade I, subclade IIa, and subclade IIb, with homologous and nonhomologous sites highlighted; (B) Entropy heatmap; (C) Entropy analysis by site.

**Figure S4. Conservation and variation in proteins encoded by orthologous poxvirus gene (OPG) 153.**

(A) Entropy heatmap. CMLV, camelpox virus; VARV, variola virus; VACV, vaccinia virus; CPXV, cowpox virus; MPXV, monkeypox virus. (B) . MPXV, monkeypox virus; CPXV, cowpox virus; VACV, vaccinia virus; VARV, variola virus; CMLV, camelpox virus.

## SUPPLEMENTAL TABLE LEGENDS

**Table S1. NovaSeq sequencing quality control values**

**Table S2. Low-complexity regions (LCRs) in the monkeypox virus (MPXV) high-quality genome (HQG) sequence 353R**

Listed are annotated positions (according to the reference MPXV-M5312_HM12_Rivers isolate genome sequence), sequence, and flanking regions of each area, as described previously (Phillips et al., 2018). ID, identification; STR, short tandem repeats.

**Table S3. Short tandem repeats (STRs) in low-complexity regions (LCRs)**

Detailed information on STRs of the entire dataset used for analysis in **Figure 2**.

**Table S4. Phylogenetic analysis**

Detailed information on groups depicted in **Figure S2**. SNP, single nucleotide polymorphism.

**Table S5. Genomes used in the study**

GI, genome identifier; NA, not applicable

**Table S6. Characterization and validation of non-randomly distributed low-complexity regions (LCRs) in the monkeypox virus (MPXV) 353R genome sequence**

Detailed information on the represented materials, along with originator and epidemiological data, used for analysis in **Figure 2**. SRA, Sequence Read Archive (SRA); ID, identification; QC, quality control.

## SUPPLEMENTAL FILE LEGENDS

**File S1. Analysis parameters for validation of short tandem repeats (STRs) in National Center for Biotechnology Information (NCBI) Sequence Read Archive (SRA)**

## Notes

### Competing Interest Statement

The work for this study at Instituto de Salud Carlos III was partially funded by Accion Estrategica Impacto clinico y microbiologico del brote por el virus de la viruela del mono en pacientes en Espana (2022): proyecto multicentrico MONKPOX-ESP22 (CIBERINFEC).
The work for this study at the GP laboratory was funded by instiutional funds of the Department of Microbiology, Icahn School of Medicine at Mount Sinai in support of Global Health Emerging Pathogen Institute activities.
The A.G.-S. laboratory has received research support from Pfizer, Senhwa Biosciences, Kenall Manufacturing, Blade Therapuetics, Avimex, Johnson & Johnson, Dynavax, 7Hills Pharma, Pharmamar, ImmunityBio, Accurius, Nanocomposix, Hexamer, N-fold LLC, Model Medicines, Atea Pharma, Applied Biological Laboratories and Merck, outside of the reported work. A.G.-S. has consulting agreements for the following companies involving cash and/or stock: Castlevax, Amovir, Vivaldi Biosciences, Contrafect, 7Hills Pharma, Avimex, Vaxalto, Pagoda, Accurius, Esperovax, Farmak, Applied Biological Laboratories, Pharmamar, Paratus, CureLab Oncology, CureLab Veterinary, Synairgen and Pfizer, outside of the reported work. A.G.-S. has been an invited speaker in meeting events organized by Seqirus, Janssen, Abbott and Astrazeneca. A.G.-S. is inventor on patents and patent applications on the use of antivirals and vaccines for the treatment and prevention of virus infections and cancer, owned by the Icahn School of Medicine at Mount Sinai, New York, outside of the reported work.

### Summary of Updates

Added Disclaimer of United States Army personnel. Corrected an error regarding the codon usage in one of the promoters of OPG204.

