## Supplementary figures and images for "Changes in a new type of genomic accordion may open the pallets to increased monkeypox transmissibility"

### Supplemental Figure 1

#### Supplementary Figure 1

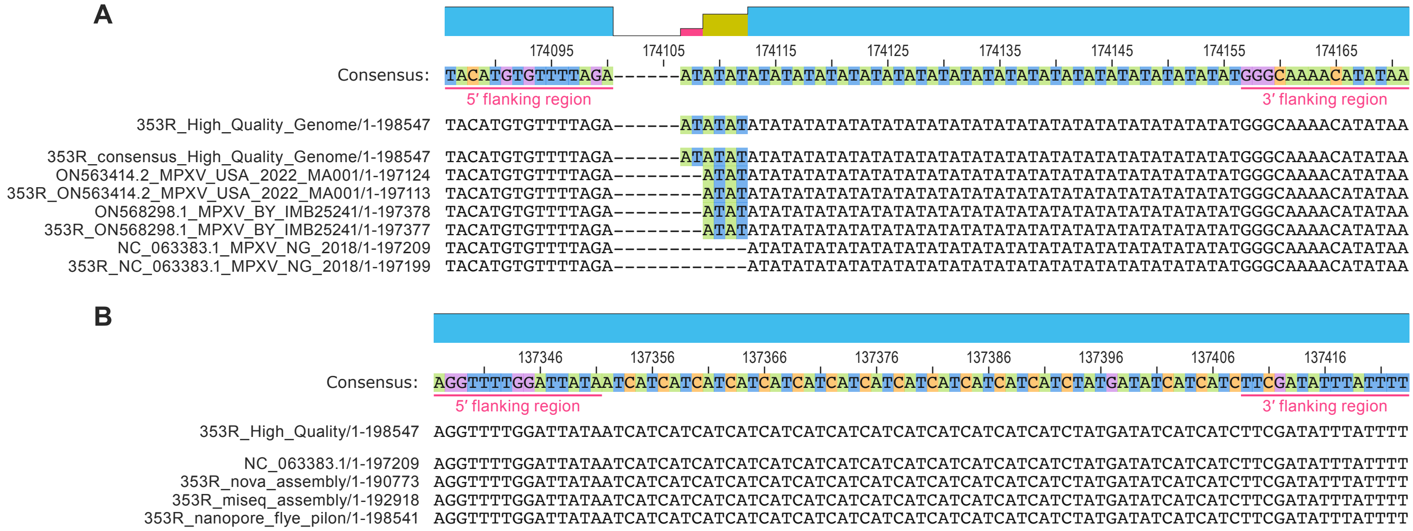

### Supplemental Figure 2

Supplementary Figure 2A

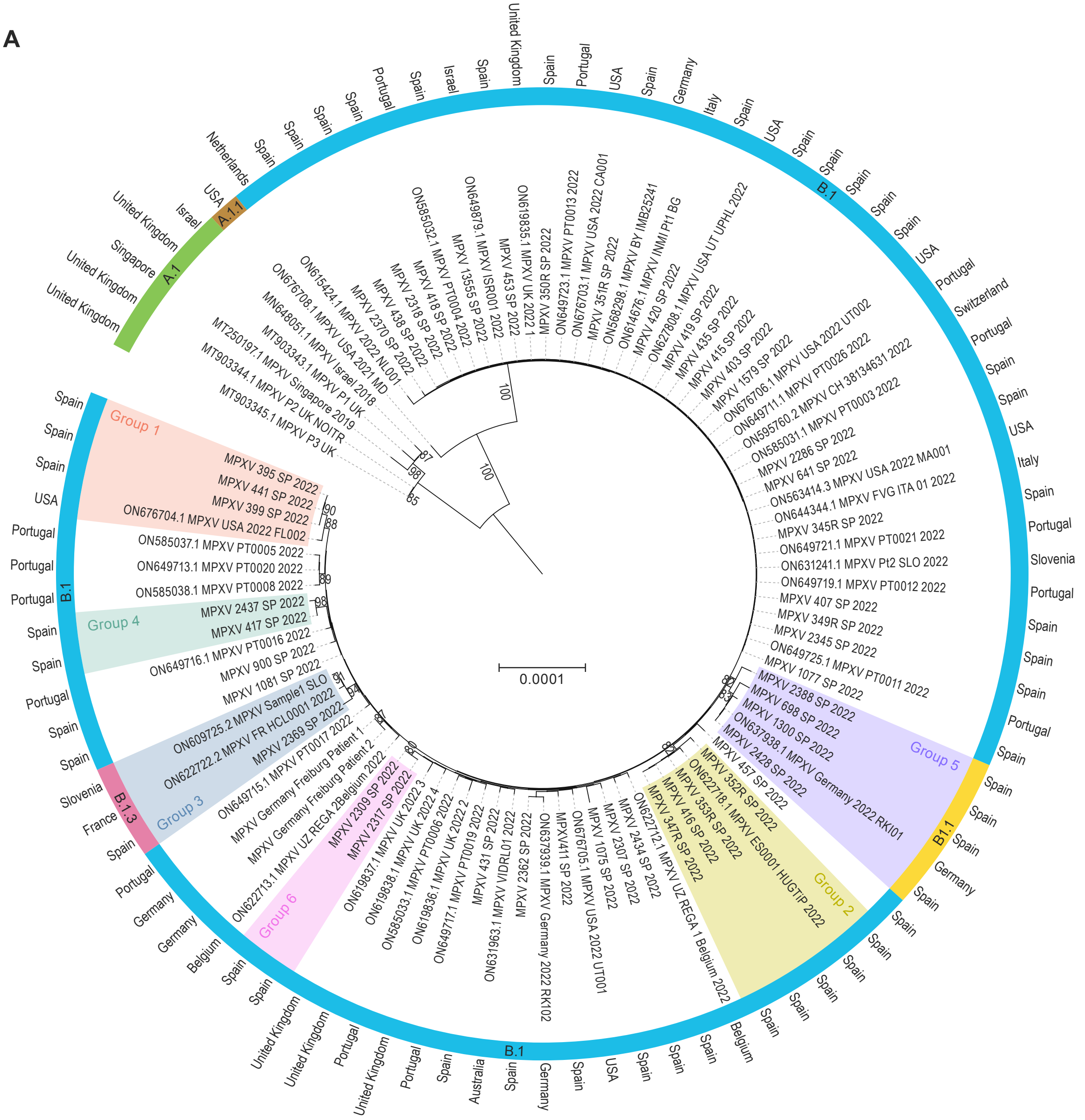

Supplementary Figure 2B

B

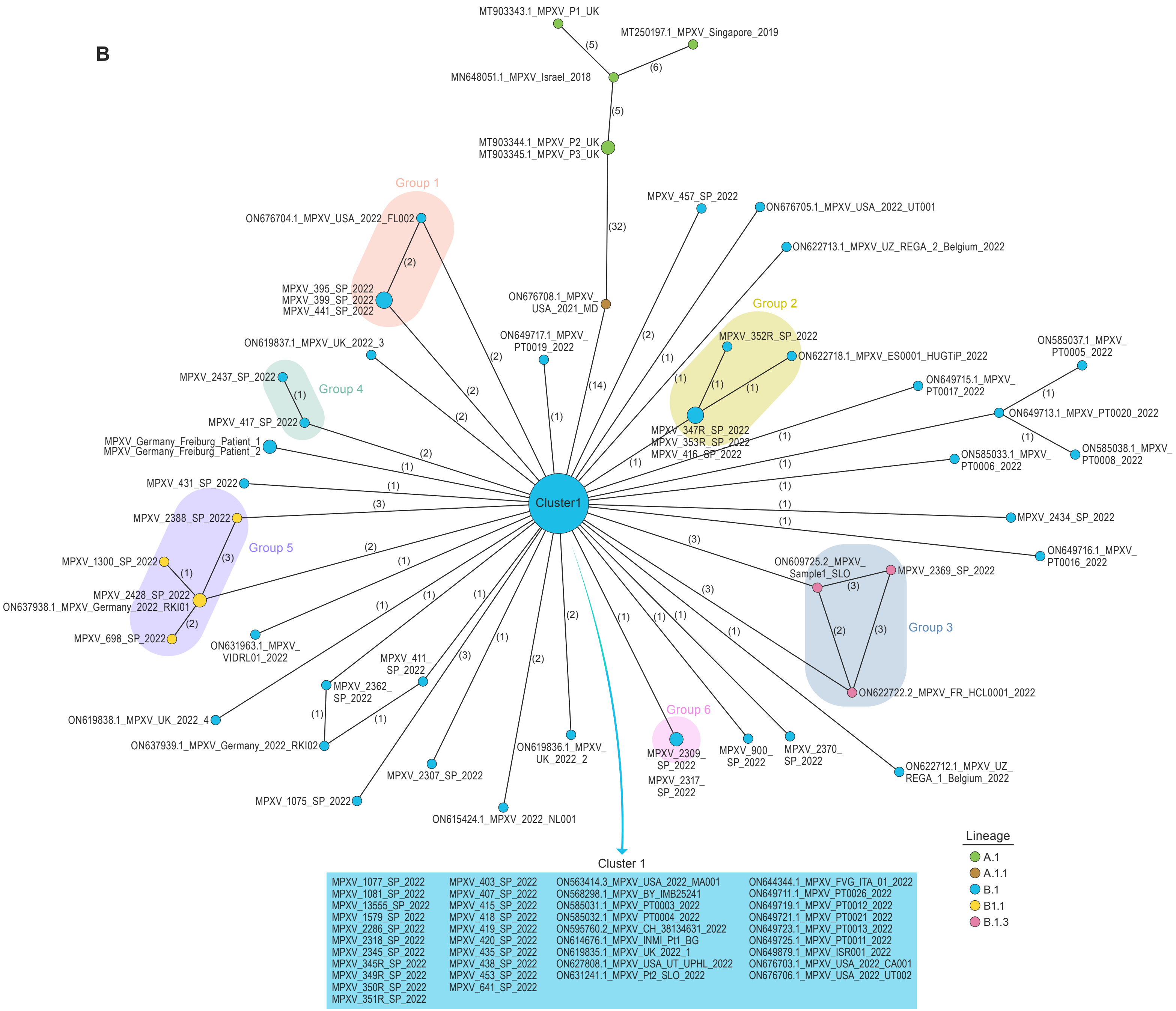

### Supplemental Figure 3

#### Supplementary Figure 3

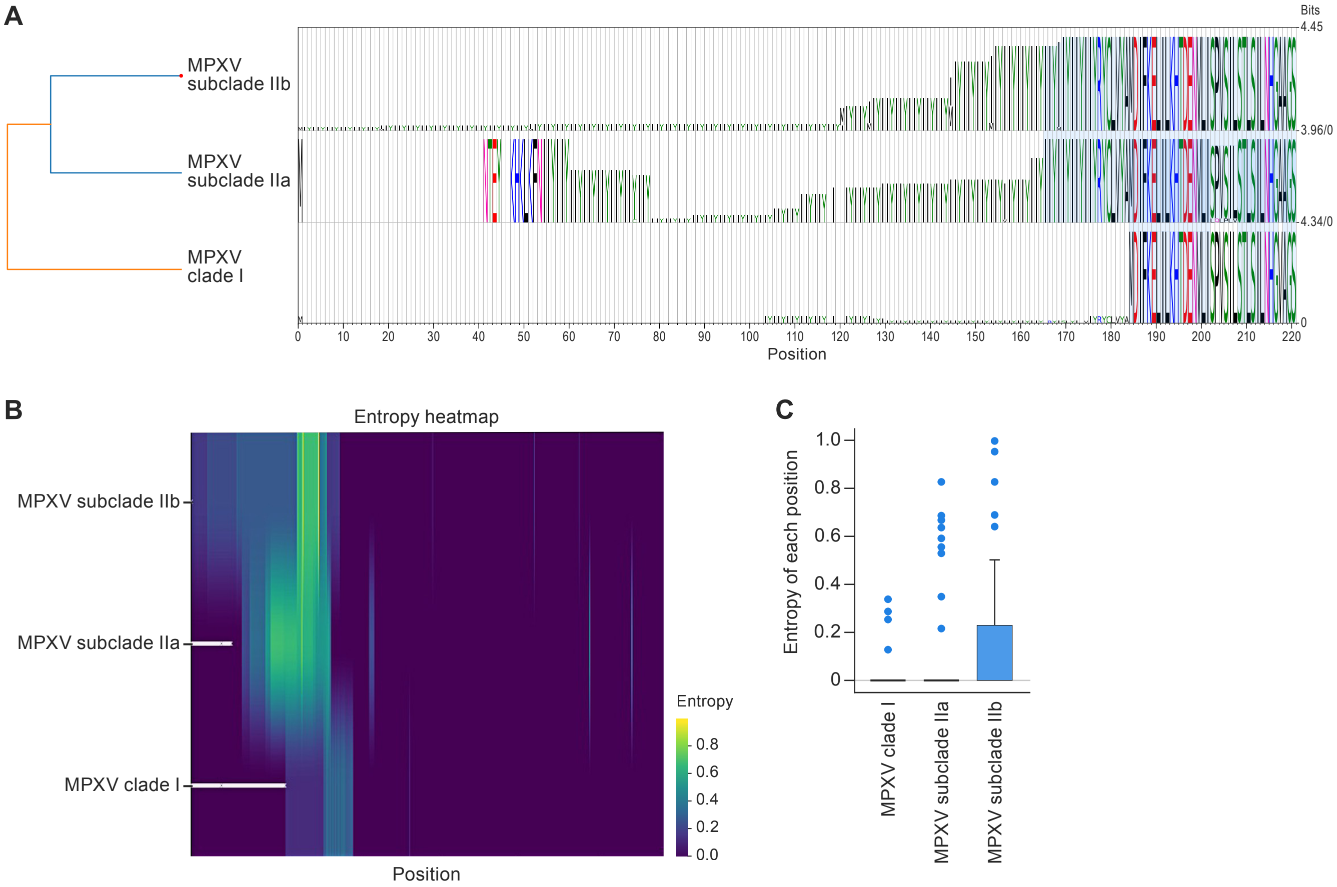

### Supplemental Figure 4

Supplementary Figure 4

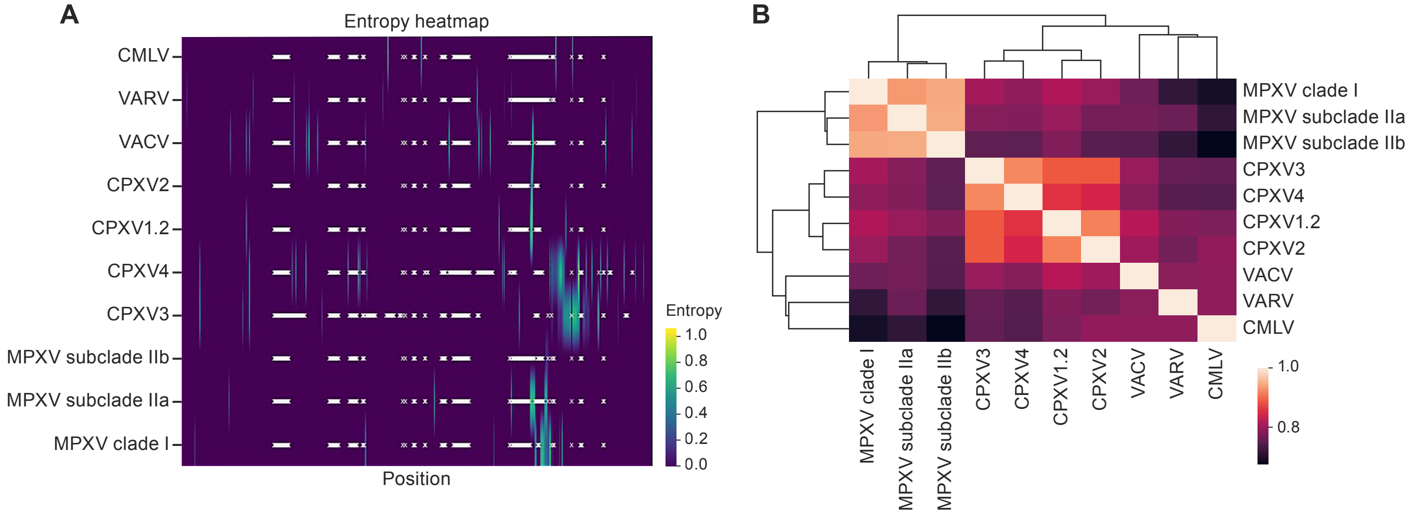
