## Supplemental File 1 for "Changes in a new type of genomic accordion may open the pallets to increased monkeypox transmissibility"

**SUPPLEMENTARY FILE 1**

A. **Short tandem repeats curation**

Short Tandem Repeats areas where manually inspected in assemblies from sample 353R. Alignments show for all identified LCRs:

1. NC_063363.1 reference genome
2. 353_R sample de novo assembly from MiSeq reads 2x 300
3. 353_R sample de novo assembly from NovaSeq 2x150 reads
4. 353_R sample de novo assembly from Nanopore reads.
5. High quality genome

**LCR1:**

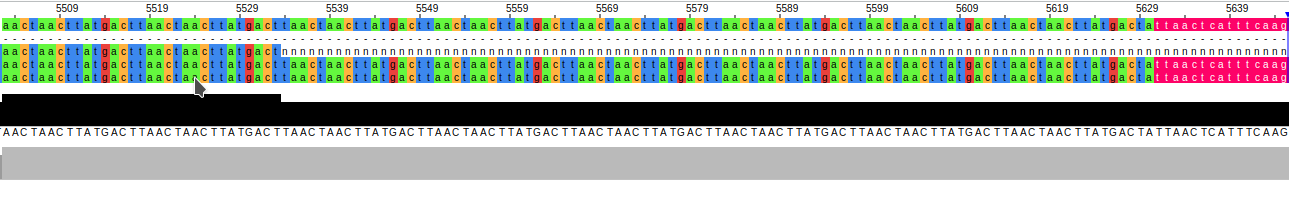

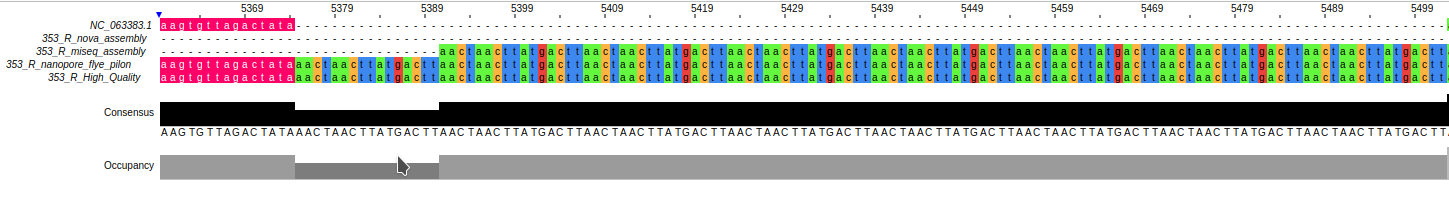

**LCR2:**

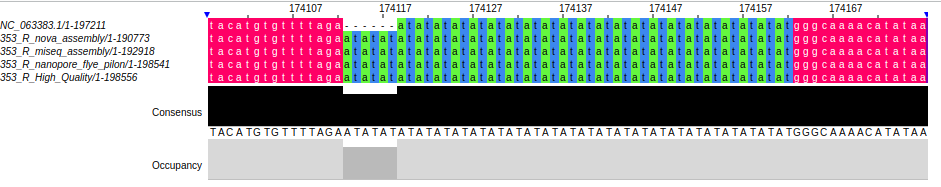

**LCR3:**

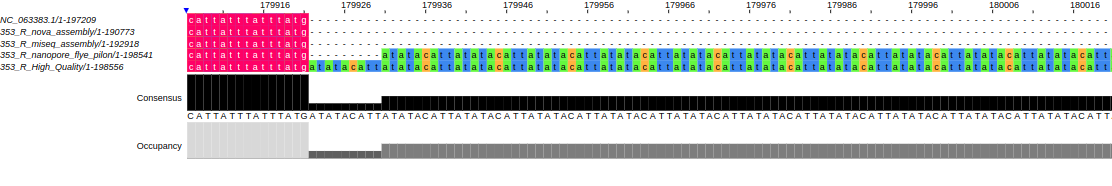

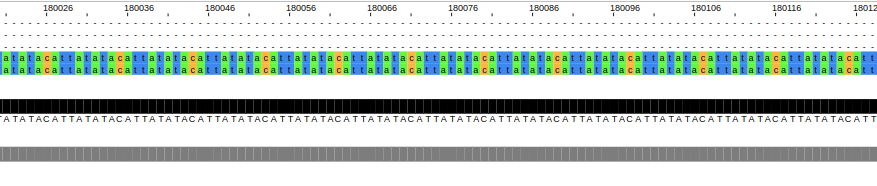

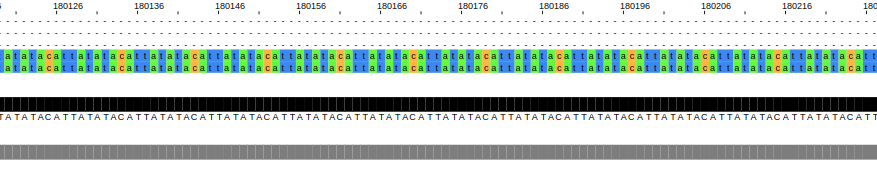

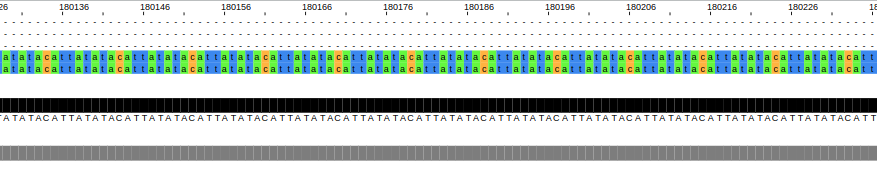

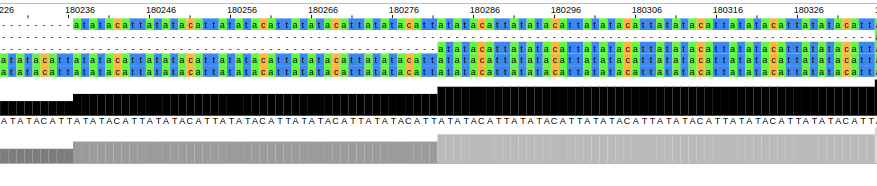

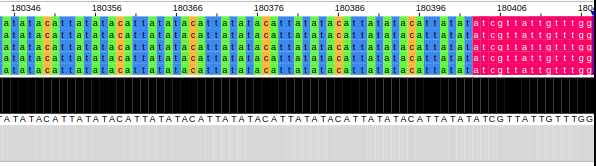

**LCR4:**

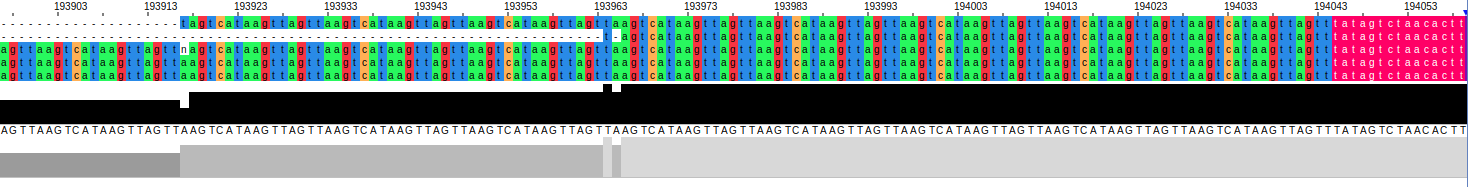

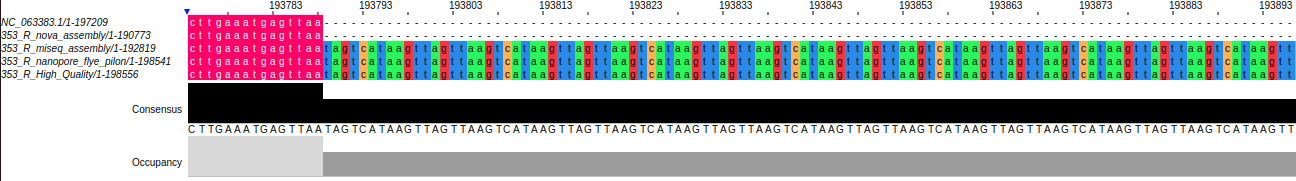

**LCR5:**

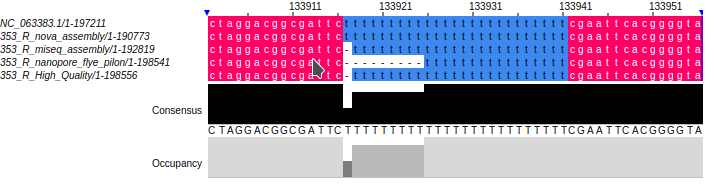

**LCR6:**

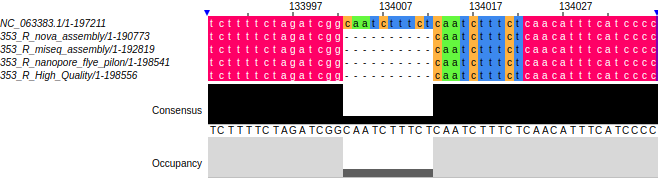

**LCR7:**

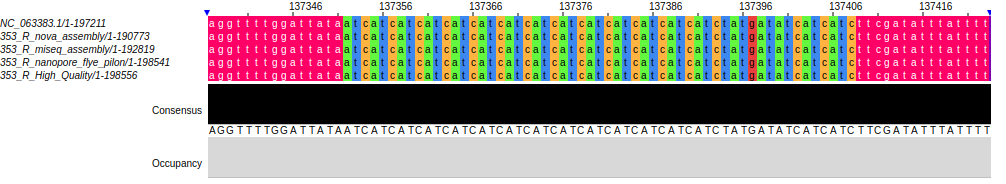

**LCR8:**

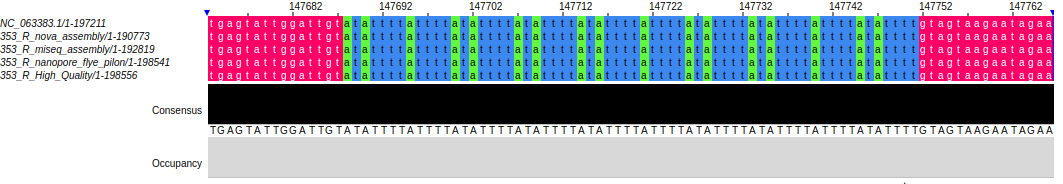

**LCR9:**

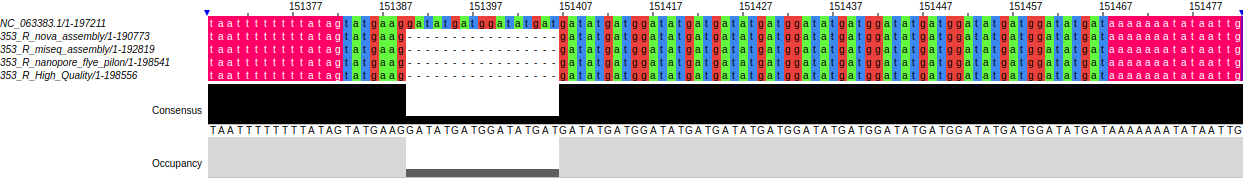

**LCR10:**

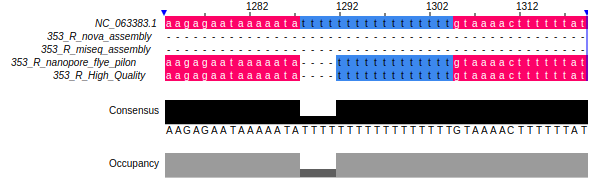

**LCR11:**

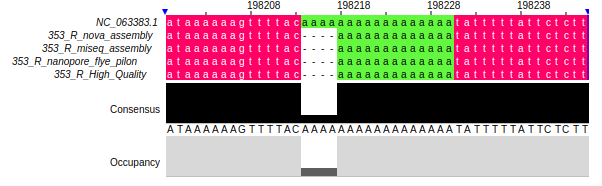

**LCR12:**

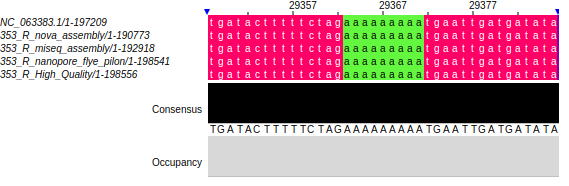

**LCR13:**

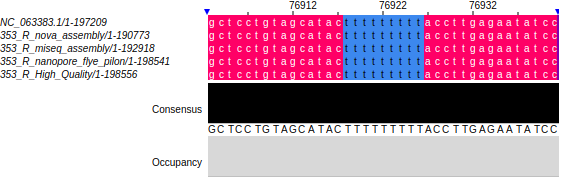

**LCR14:**

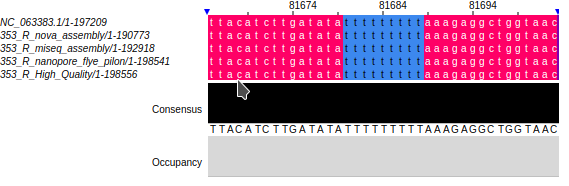

**LCR15:**

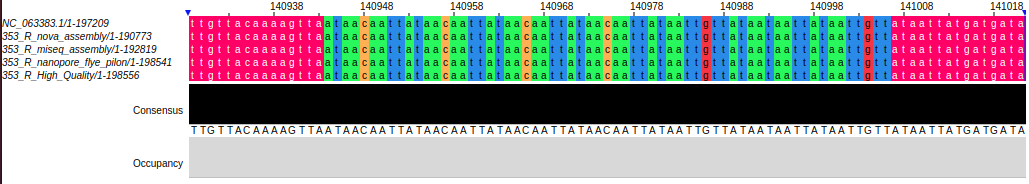

**LCR16:**

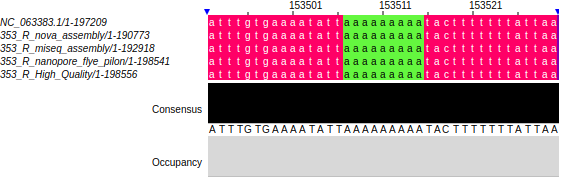

**LCR17:**

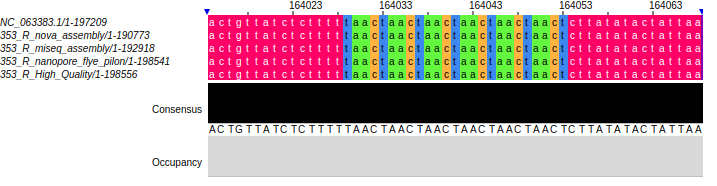

**LCR18:**

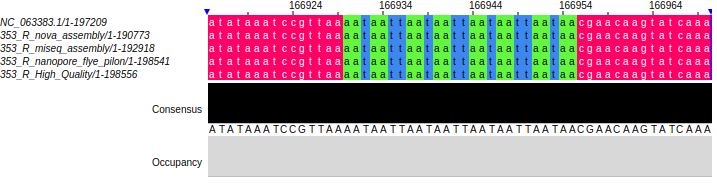

**LCR19:**

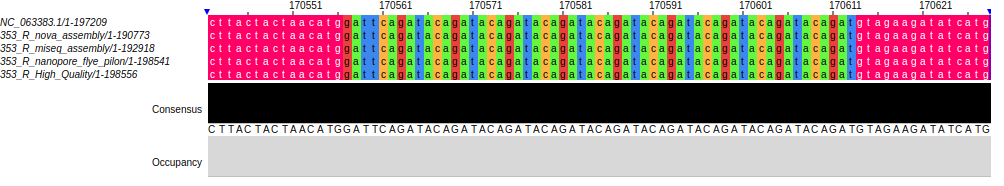

**LCR20:**

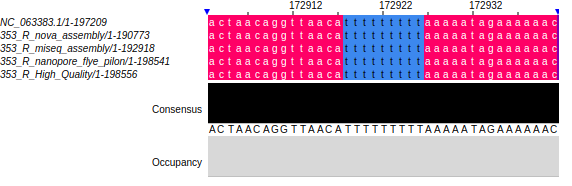

**LCR21:**

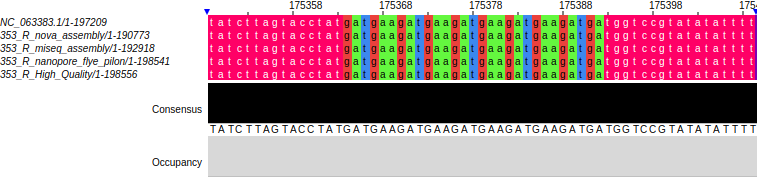

**B. Nanopore samples analysis downloaded from SRA**

Monkeypox samples with nanopore data available in SRA (August 10, 2022) were downloaded from NCBI webpage (<https://www.ncbi.nlm.nih.gov/sra>) resulting in 35 samples. Next, reads were de novo assembled using flye assembler v2.9-b1768 [(Kolmogorov et al. 2019)](https://sciwheel.com/work/citation?ids=6744810&pre=&suf=&sa=0) with default parameters in nanopore raw reads mode, and analyzed using a modified version of strsearch (<https://github.com/BU-ISCIII/MPXstreveal>). Reads spanning the defined LCR3, LCR1 and LCR4 regions were identified. Those with both flanking reads with 0 mismatches were collected, aligned and generated a consensus sequence. Potential number of repeats of selected LCRs in each sample were inspected comparing the assembled genome and the strsearch result according to **Supplementary table 6**.

The “de novo'' assembly method only resulted in the resolution of a few complete genomes. To identify the number of LCRs repeats, we mostly used the method spanning the flanking regions. Although the results differ in some cases between bioinformatic methods, and we cannot validate the exact numbers of repeats experimentally since we do not have access to the samples, the values obtained show clear differences in the tendency of number of repeats. For clarity, the most conservative number was selected, considering the biology of MPXV replciation, the solution that was supported by the highest number of genomic data, and the selection of the solution that underestimated variation.
